# Psychedelics restructure information in the human brain

**DOI:** 10.64898/2026.10.01.755915

**Authors:** George Blackburne, Alberto Liardi, Pablo Mallaroni, Rosalind G. McAlpine, Robin L. Carhart-Harris, Christopher Timmermann, Suresh Muthukumaraswamy, Jeremy I. Skipper, Pedro A. M. Mediano

## Abstract

How do psychedelic drugs alter human brain function? Answering this question relies on our ability to understand how information is stored, transferred, and transformed across the cortex. Here, we leverage a precise mathematical framework for multivariate information decomposition to construct the first comprehensive picture of these processes across five compounds: LSD, psilocybin, ketamine, DMT and 5-MeO-DMT (N=80). We provide a formal and empirical explanation of major prior results, revealing sets of competing underlying phenomena masked by extant approaches. We extend this analysis to novel higher-order phenomena, and we demonstrate how all of these observations can emerge from a minimal generative model in which neural populations operate with reduced self-coupling. In sum, our results suggest that the psychedelic brain is one in which cortical populations become alike, but not necessarily together. This has important implications for models of how psychedelics reorganize brain dynamics to alter subjective experience and alleviate mental health conditions.

## I. INTRODUCTION

In the last decade, scientific interest in psychedelic drugs has resurged, fueled by their potential both as experimental tools for understanding how the brain orchestrates exotic mental phenomena and as therapeutic agents for treating mental health conditions [1, 2]. This renewed focus has entailed a rapid expansion in the collection and analysis of neuroimaging data of humans under the influence of various psychedelics [3–5]. However, at present, the neuroscientific study of psychedelics suffers from a lack of empirical unification, that is, the joint investigation of multiple datasets for multiple drugs under a comprehensive mathematical framework. This likely reflects not only the characteristic structural fragmentation of an emerging scientific field, but also the non-trivial task of assembling an exhaustive enough framework to accommodate the multitude of ways brain activity might express itself.

This lack of unification is perhaps ironic, given that the working hypothesis across models of psychedelic action, whether explicit or implicit, is that psychedelics increase “integration” in the brain [5]. Contemporary work routinely tells the story that psychedelics “liberate” [6] neural dynamics from the functional segregation that defines putative default circuits and systems [7], fostering instead a more collective information sharing regime marked by global connectivity [2, 5, 8–12]. Additionally, some theories place significant explanatory weight on the idea that, concomitant with this increased integration of neural dynamics across space, there is a rise in the local differentiation of neural dynamics across time, often framed as psychedelics augmenting the level of “complexity” of brain dynamics [13]. Indeed, these appear so uncontroversial that the only point of deliberation remaining is what neurobiological process might underlie them, be it a breakdown in thalamo-cortical gating [14], the claustro-cortical coordination that entrains modular brain function [15], or the precision weighting that governs stern predictive coding hierarchies [16]. In other words, we argue over the neurobiological explanans of a behaviour that we have potentially prematurely converged on as the crucial explanandum.

While this foundational work has been crucial, the overlooked question is: what exactly do we talk about when we talk about the integration of information? Does it mean regions simply holding the same information, information moving from one region to another, or information being jointly modified by those regions? More pragmatically, when researchers discuss integration in the context of functional connectivity (the co-fluctuation of regions measured by Pearson’s correlation), do they mean the same thing as when discussing directed functional connectivity (the predictive influence of one region on another measured by Granger Causality)? These questions illustrate that “integration”, “interaction”, “information sharing”, “communication” and “connectivity” are often underspecified concepts in neuroscience, vague objects of study further obscured by our ways of measuring them. Making progress on this problem therefore, relies on our ability to recognize and formalize the diversity of ways information may be integrated, or differentiated, across space and time, in a complex dynamical system like the brain.

Recent advancements in information theory have begun to address this problem, with the development of information decomposition approaches that can reveal the fundamental “atoms” by which information can be stored, transferred, or transformed [17–20]. Importantly, recent work in this field has shown both that the standard measures in our scientific toolkit are systematically blind to particular modes of information, and that quantities designed to capture distinct phenomena in fact detect overlapping informational atoms [20]. Thus, with the computational and conceptual landscape made available by the grammar of information dynamics, we are able to concretely ask: where is information coming from, and where is it going in the psychedelic brain?

Here, we introduce Integrated Information Decomposition (ΦID) as a formal framework for the comprehensive decomposition of information dynamics in neural activity [20], and present a vocabulary for its derived quantities. We then employ this to explain the informational basis of the main prior results in the literature on the effects of psychedelics (LSD, psilocybin, ketamine, DMT, and 5-MeO-DMT) on the human brain under the same mathematical roof, and extend these insights with novel measures that capture previously undetectable phenomena. Finally, we provide a generative computational model that reproduces the effects of psychedelics on information dynamics. In doing so we yield a comprehensive, unified, and mechanistic lens on the large-scale brain action of psychedelic drugs.

## II. RESULTS

### A. A unified taxonomy of information dynamics

To exhaustively identify the modes of information dynamics executed within and across brain regions over time, we employ Integrated Information Decomposition (ΦID) (see Methods) [20]. Treating the past activity of each brain region as sources 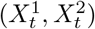, and the future of those same regions as targets 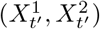, ΦID performs a decomposition of the information flow between them. Like other decomposition approaches [17–19], ΦID identifies three flavors of information at each instance: *redundancy*, which represents information that is identically held or repeated over each element, *synergy*, which represents the information that is contained only in the joint but not the marginal state of the elements, and *unique* information, which represents the information that is provided by one and only one particular element. In contrast to previous decomposition frameworks, in which these forms of information are defined with respect to a particular time coordinate, ΦID enables the characterization of the structure of information with respect to the past and the future, and thus how information can stay or change format over time. For example, a bit of information that was previously held redundantly in the past may become uniquely held by a particular element in the future [20]. In the following, we introduce the atomic quantities of ΦID and the kind of information dynamics they capture.

To understand these modes of information, we introduce some nomenclature (Fig. 1). If information is redundant in the past and remains redundant in the future, we denote this as *Oneness* [21], since information is identically represented at every spatial and temporal coordinate. If information is common among the elements in the past, but comes to be held by one specific element in the future, we denote this as *Erasure*, since it has been removed from the other element over time. The time-reversed counterpart of Erasure, *Copy*, is therefore information which is contained in one element in the past but in all elements in the future. In other words, it indicates that the information has replicated over time. If information is uniquely held by a particular element, and is stored exclusively there over time, we term this *Preservation*. Alternatively, if information is uniquely held by a particular element, and is then transferred such that it becomes unique to the other, we term this *Migration* [22].

**FIG. 1.**
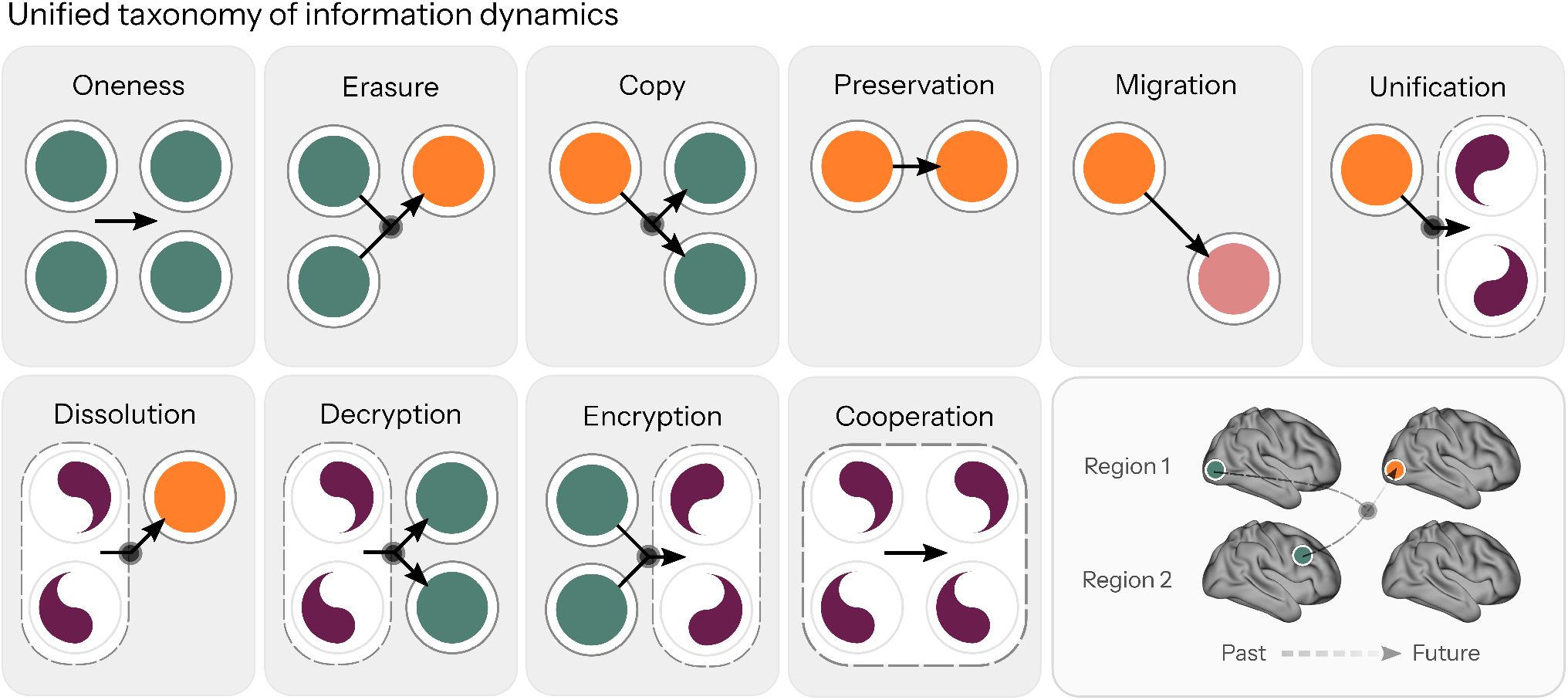
A unified taxonomy of information dynamics. Comprehensive decomposition of the ways information is stored or transferred in and between the parts, wholes, past, and future of the brain via Integrated Information Decomposition (ΦID) [20]. Specifically, this details how information can remain, or be exchanged, over time via three distinct formats: Redundant (Green) information is held identically by both brain regions simultaneously, Synergistic (Purple) information is held exclusively in the joint state of both brain regions, and Unique (Pink or orange) information that is held by one brain region and that region only. Maintenance or transformation between these formats enables the possibility of ten qualitatively distinct modes of information dynamics (from left to right): Oneness, Erasure, Copy, Preservation, Migration, Unification, Dissolution, Decryption, Encryption and Cooperation.

The notions introduced thus far have only encompassed redundant and unique information. However, arguably the most interesting quantities are those that involve synergistic information. If information is uniquely held by one element in the past, but becomes specified in the joint state of the elements in the future, we term this *Unification*. Its time-reversed counterpart, *Dissolution*, is when information is integrated in the whole in the past but becomes disintegrated and held independently by one element in the future. Similarly, *Decryption* and *Encryption* [22] refer to synergistic information in the past becoming redundant information in the future, and vice versa, representing the transformation of information between higher-order de-localized and lower-order multi-localized formats. Finally, information coded in the whole persistently both in the past and in the future, we term *Cooperation* [21], as this captures information that emerges via the dynamical complementarity of the parts that transcends their individual contributions. With this terminology, we provide an intuitive process-based lexicon for interpreting the information dynamics revealed by ΦID.

With this formalism and nomenclature in hand, we return to the present question: How should we understand the effects of psychedelic drugs on large-scale human brain function? To answer this, we apply ΦID to the evolving interaction between each and every brain region, from previously collected source-localized parcellated M/EEG data from five studies of healthy human subjects under the influence of classical and atypical psychedelic drugs – including lysergic acid diethylamide (LSD), 4-phosphoryloxy-N,N-dimethyltryptamine (psilocybin), ketamine, N,N-dimethyltryptamine (DMT), and 5-methoxy-N,N-dimethyltryptamine (5-MeO-DMT).

### B. Explaining existing measures in terms of information

We begin by establishing contact with the prior literature in asking: what is the informational basis of the main contemporary empirical results on the effects of psychedelics on neural activity?

#### 1. Complexity

One of the most consistently reported effects of psychedelic drugs is a global increase in neural signal diversity or “complexity” [12, 23–33] (but see [34]). The vast majority of studies have investigated this effect through approximations of the entropy rate 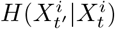, using estimators such as Lempel-Ziv Complexity [31, 35]. Entropy rate quantifies the “complexity” or “innovation” of a time-evolving process as a departure from the “inheritance” of information transmitted from the past to the future 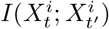, a quantity also known as the active information storage (AIS). More formally, we have:

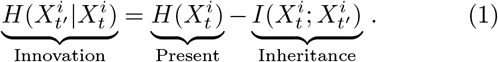

This relationship is simple: the less captive a stochastic neural process is to its history, the freer it is to invent.

Indeed, empirically, this dependency is clear: across all five drugs, changes in complexity via state-space entropy rate [31] strongly anti-correlate with changes in AIS (min *r* = *−*0.88 5-MeO-DMT, max *r* = *−*0.91 LSD, PSIL, KET, Fig. 2a), as can also be seen viewing each condition in isolation (Supplementary Fig. S1). Importantly, though, the AIS of a particular brain region will not just capture information that is unique to that brain region, but also information that is redundant across other brain regions. Indeed, ΦID reveals that the AIS captures the sum of multiple atoms of information:

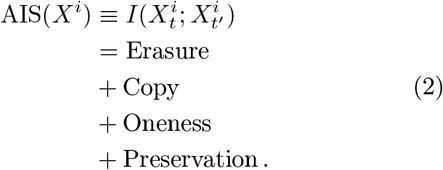

**FIG. 2.**
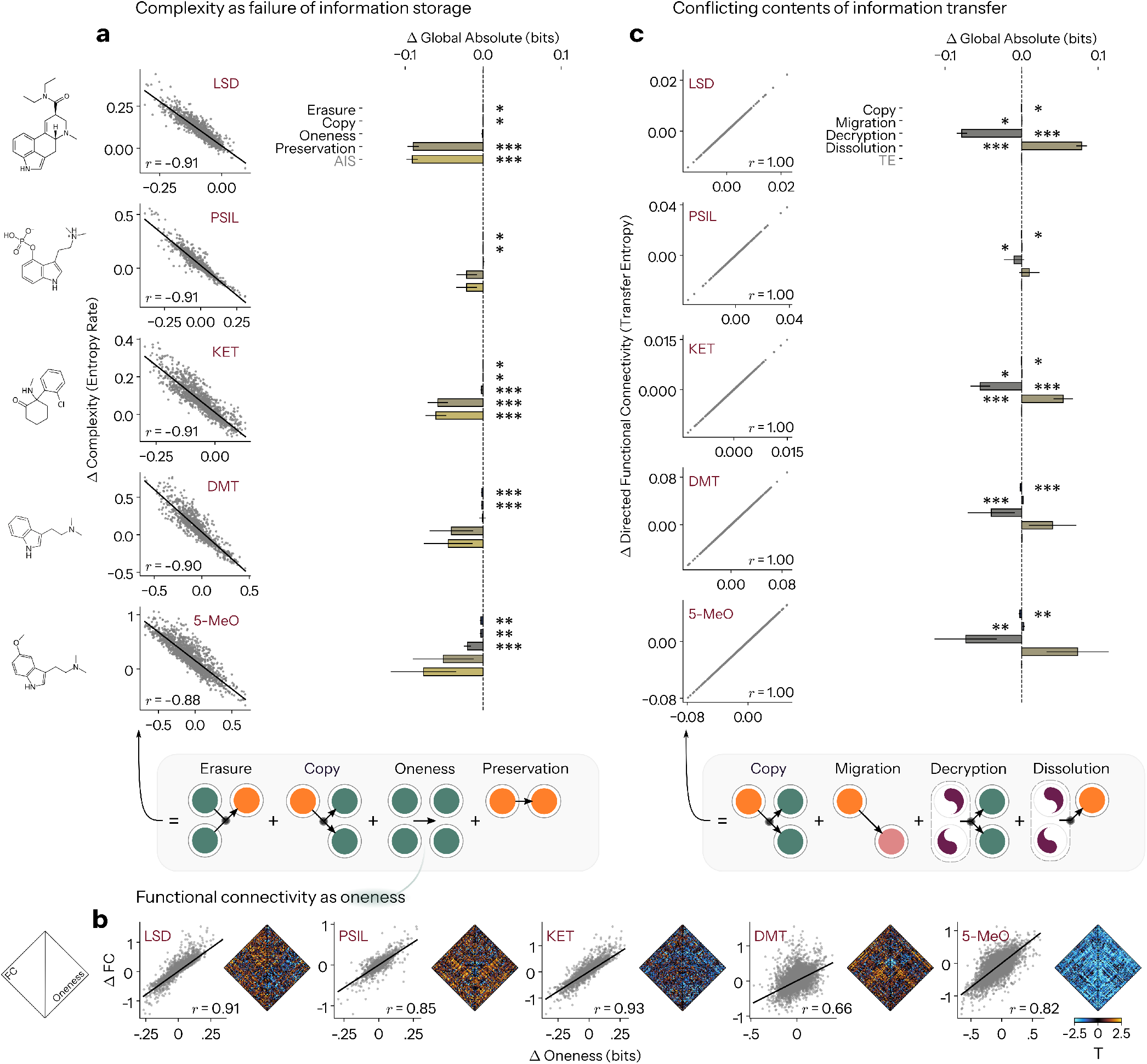
The information dynamical nature of prior findings. **a** Complexity via entropy rate depends on loss of active information storage (AIS) (left column). The AIS is itself an aggregate quantity, being the sum of four modes of information dynamics: Erasure, Copy, Oneness, and Preservation (bottom). Global absolute changes in AIS, Erasure, Copy, Oneness, and Preservation across all five drugs. Psychedelics induce heterogeneous effects on each mode of information dynamics, while Preservation is the fundamental driving factor of AIS. **b** Functional connectivity (FC) depends on Oneness, a form of information dynamics also detected by the AIS. Statistical differences in FC (left triangle) almost exactly mirror Oneness (right triangle). **c** Directed functional connectivity via Transfer Entropy (TE) is the sum of four modes of information dynamics: Copy, Migration, Decryption and Dissolution. Copy is therefore detected by both AIS and TE. Global absolute changes in TE, Copy, Migration, Decryption and Dissolution across all five drugs. Psychedelics again induce heterogeneous effects on each mode of information dynamics, with Decryption and Dissolution showing competing behaviors. (*, **, *** indicate *p <* .05, .01, .001, FDR-corrected Benjamini-Hochberg procedure. Bars indicate mean *±* SEM. psilocybin (PSIL), ketamine (KET), 5-MeO-DMT (5-MeO)).

That is, the AIS captures the sum of Erasure, Copy, Oneness, and Preservation. These four quantities can vary independently of each other, and it is trivial to construct systems with the same AIS with entirely different atomic contributions. Crucially, these distinct processes carry different interpretations, meaning it is necessary to ask what drives global changes in AIS. While we find that loss of Preservation is generally the strongest numerical factor, showing global mean reductions across all drugs, this is only significant for LSD and ketamine (LSD: *p*_FDR_ = 1.25 *×*10^*−*8^, PSIL: *p*_FDR_ = 0.194, KET: *p*_FDR_ = 4.92*×* 10^*−*4^, DMT: *p*_FDR_ = 0.200, 5-MeO: *p*_FDR_ = 0.202). On the other hand, we find that Erasure and Copy are significantly reduced across all five compounds (Erasure: LSD *p*_FDR_ = 0.039, PSIL *p*_FDR_ = 0.018, KET *p*_FDR_ = 0.012, DMT *p*_FDR_ = 2.0 *×*10^*−*4^, 5-MeO *p*_FDR_ = 0.008; Copy: LSD *p*_FDR_ = 0.039, PSIL *p*_FDR_ = 0.016, KET *p*_FDR_ = 0.012, DMT *p*_FDR_ = 2.1 *×*10^*−*4^, 5-MeO *p*_FDR_ = 0.008, respectively; regional results in Supplementary Fig. S4). Finally, ketamine and 5-MeO-DMT both exhibit significant reductions in Oneness (KET: *p*_FDR_ = 6.82 *×* 10^*−*4^, 5-MeO: *p*_FDR_ = 5.4 *×* 10^*−*4^).

#### 2. Functional connectivity

Another consistently reported effect of psychedelic drugs is alterations in Pearson’s correlation (*ρ*), otherwise known as “functional connectivity” (FC), across the human cortex [8]. Naturally, we then ask what type of information dynamics this reflects. FC cannot be exactly derived from the present framework, as it is a statistic that is blind to the passage of time, i.e. it relies exclusively upon the steady-state distribution of the data. However, a reasonable approximation can be derived in terms of Oneness. Under Gaussian assumptions, the instantaneous mutual information *I*(*X*^*i*^; *X*^*j*^) between two regions can be obtained from FC as:

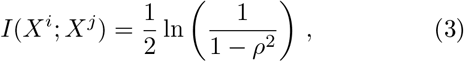

where *ρ* is the correlation between *X*^*i*^ and *X*^*j*^. Moreover, in the present work, Oneness is defined via the minimum mutual information within and between regions:

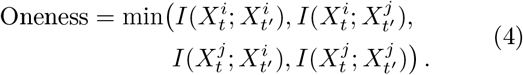

Given that likely universally in real-world dynamical systems, including empirical neuroimaging data, the lagged cross-correlation is lower than the autocorrelation 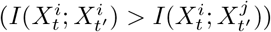, and the time delayed mutual information is upper bounded by the instantaneous mutual information 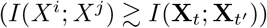. We can then put everything together and write:

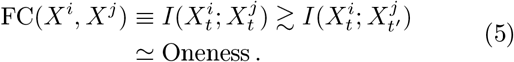

Indeed, we find that changes in Oneness strongly correlate with changes in FC (min *r* = 0.66 DMT, max *r* = 0.93 KET) (Fig. 2b). This result is consistent with prior work that shows a strong association between FC and redundancy, across different decompositions [36, 37].

A central result here is that, viewed this way, FC is an element detected in the AIS, and as just discussed, therefore constrains entropy rate (Fig. 2a).

#### 3. Directed functional connectivity

Finally, we turn to directed functional connectivity (dFC). In previous studies, dFC has been assessed via Granger Causality [25, 38] and Transfer Entropy (TE) [39], quantities that are equivalent for Gaussian variables [40]. However, like for AIS, ΦID reveals that the TE 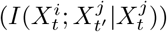 captures the sum of multiple atoms of information:

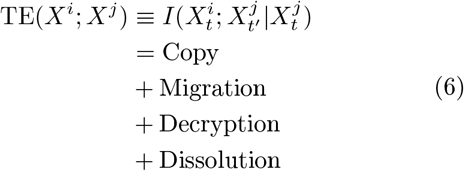

Importantly, like AIS, one can see that the TE includes Copy, showing that classical measures of information storage and information transfer, previously assumed to be independent, in fact measure overlapping phenomena [20].

Examining global changes in each atom, we find a competitive effect of Decryption and Dissolution across drugs: psychedelics tend to decrease mean levels of global Decryption while concomitantly increasing Dissolution, though this effect varied considerably in magnitude and significance across substances. LSD and ketamine showed robust effects (LSD: *p*_FDR_ = 3.57*×* 10^*−*8^, *p*_FDR_ = 3.47*×* 10^*−*8^; KET: *p*_FDR_ = 7.38*×* 10^*−*4^, *p*_FDR_ = 7.35 *×*10^*−*4^), whereas psilocybin, DMT, and 5-MeO-DMT did not reach significance (PSIL: *p*_FDR_ = 0.440, *p*_FDR_ = 0.437; DMT: *p*_FDR_ = 0.263, *p*_FDR_ = 0.262; 5-MeO-DMT: *p*_FDR_ = 0.143, *p*_FDR_ = 0.141; for Dissolution and Decryption respectively throughout). Interestingly, Migration significantly increases under all five drugs, most strongly for DMT (LSD: *p*_FDR_ = 0.039; PSIL: *p*_FDR_ = 0.016; KET: *p*_FDR_ = 0.012; 5-MeO-DMT: *p*_FDR_ = 0.008; DMT: *p*_FDR_ = 2.0 *×*10^*−*4^). Thus, psychedelics exert coherent and differentiable effects on the underlying constituents of canonical formalisms of information transfer.

In summary, these results elucidate the informational structure of the main quantities utilized in the neuroimaging, particularly with respect to psychedelics: signal complexity, functional connectivity, and directed functional connectivity. These results show how classic measures assumed to assess orthogonal properties measure, in fact, overlapping ones, and how such measures can obscure competing effects occurring under the hood. Importantly, analysing the basic information dynamics on their own terms reveals a suite of convergent and divergent behaviors across different psychedelic drugs.

### C. Reconfigurations of relative information in the psychedelic brain

While the previous section explains the informational origin of existing measures, to properly understand the effects of psychedelics on basic modes of information dynamics, it is necessary to establish appropriate controls via null models. This is critical for the information decomposition approach presented here, as it operates by partitioning the Time-Delayed Mutual Information (TDMI, 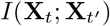. Thus, comparing the atoms of two different conditions may yield results purely dictated by differences in their mutual information. Put simply, a system with a lower value of TDMI will bias atoms to be smaller than the same atom belonging to a system with a larger TDMI. Indeed, we find that psychedelics reduce the TDMI (LSD: *p*_FDR_ = 1.07*×* 10^*−*8^, psilocybin: *p*_FDR_ = 0.155, ketamine: *p*_FDR_ = 3.99*×* 10^*−*4^, DMT: *p*_FDR_ = 0.169, 5-MeO-DMT: *p*_FDR_ = 0.151, Supplementary Fig. S2), as hinted by the finding that they increase entropy rate. To account for this, we develop a null model procedure for ΦID that builds on the approach introduced by Ref. [41] for PID (see Methods). We therefore make a distinction between absolute (unnormalized) quantities and relative (normalized) quantities: the former tells us the overall magnitude of each kind of information in the system, whereas the latter portrays the potency of a particular form of information dynamics with respect to Gaussian systems with equivalent TDMI.

#### 1. Existing information dynamics

First, we present relative alterations in the information quantities to which existing measures are sensitive: Erasure, Copy, Oneness, Preservation, Migration, Decryption and Dissolution (Fig. 3a).

**FIG. 3.**
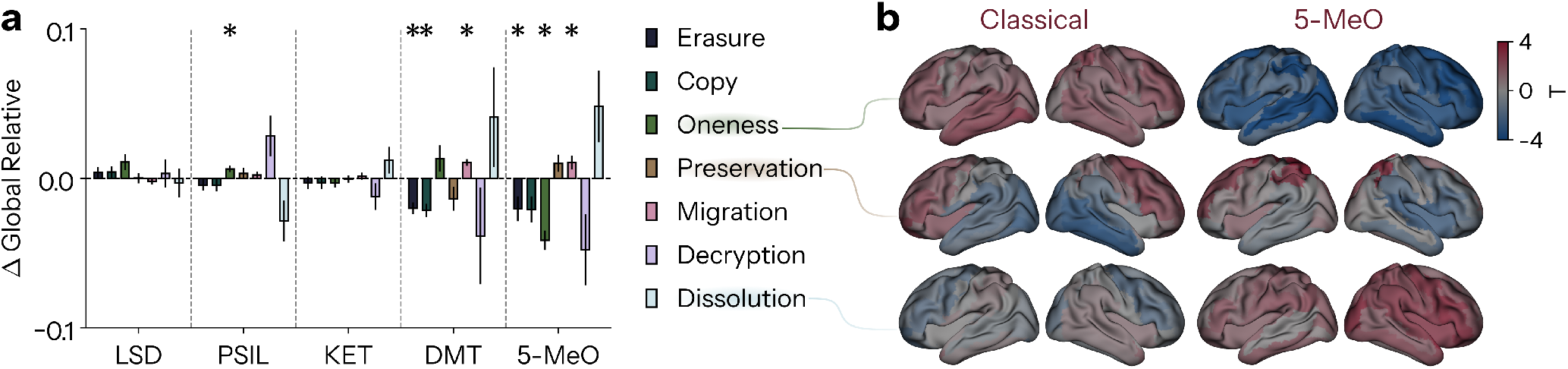
Relative reconfigurations in information dynamics reveal opposing effects of classical psychedelics versus 5-MeO-DMT. **a** Global relative changes in Erasure, Copy, Oneness, Preservation, Migration, Decryption and Dissolution. **b** Topographic statistical maps for regional differences in relative Oneness (first row), Preservation (second row), and Dissolution (third row) averaged across the classical serotonergic psychedelic drugs LSD, psilocybin and DMT (first column) and for 5-MeO-DMT (second column). Red indicates increases under drug, and blue indicates reductions. (* indicates *p <* .05, FDR-corrected Benjamini-Hochberg procedure. Bars indicate mean *±* SEM.)

We find that while classical psychedelics (LSD, psilocybin, and DMT) tend to increase relative Oneness across cortex, 5-MeO-DMT reduces it with equivalent diffusivity, with particularly impressive effects anchored in right middle frontal gyrus (BF_10_ = 3820) and right anterior cingulate (BF_10_ = 3028). Fascinatingly though, for Preservation under classical psychedelics, instead of finding homogeneous decreases across cortex, we observe a clear anterior-posterior gradient in which occipital and temporal regions decrease, and frontal regions increase. These frontal increases are remarkably strong in some instances, with right superior frontal gyrus increasing under psilocybin with BF_10_ = 838, and right mid-orbital frontal gyrus with BF_10_ = 139 (Fig. 3a,b). 5-MeO-DMT, on the other hand, shows no such gradient. Instead, it produces the strongest relative increases in Preservation of any drug (global: *T* = 4.02, *p*_FDR_ = 0.004), concentrated in medial sensorimotor and parietal cortex, peaking in right paracentral lobule (BF_10_ = 4.04 *×*10^3^) and left supplementary motor area (BF_10_ = 1.09*×* 10^3^). Our relative results reveal additional intricacies. Under DMT, Erasure and Copy are reduced most strongly in bilateral posterior cingulate (left: BF_10_ = 5.04*×* 10^3^ and 3.85*×* 10^3^ respectively), with further reductions in bilateral middle occipital gyrus (left: BF_10_ = 17.7 and 17.3). 5-MeO-DMT similarly reduces Erasure and Copy in bilateral middle occipital gyrus (right: BF_10_ = 165 and 160), but only weakly in left posterior cingulate (BF_10_ = 2.77 and 2.69) (Supplementary Fig. S6). Intriguingly, we find that 5-MeO-DMT shows the strongest mean increases in global relative Dissolution, and decreases in Decryption (Fig. 3a), potentially paralleling the global mode of deconstructed consciousness induced by the drug [42, 43], although this result is non-significant (Dissolution: *T* = 2.05, *p*_FDR_ = 0.138, Decryption: *T* = *−*2.07, *p*_FDR_ = 0.134).

#### 2. Creating and maintaining synergistic information

As previously noted, existing measures are not just unable to identify which particular form of information dynamics drives their behavior, but are also systematically blind to several other modes of information dynamics, namely: Encryption, Unification, and Cooperation (Fig. 4a). These unexplored modes involve the maintenance or dynamical creation of synergistic information, that is, information that reflects combination of multiple sources at the macroscale, and are therefore more aligned with the colloquial understanding of what it means to “integrate information”. Thus, if psychedelics work by increasing “integration” in the brain, this is where we should expect to see it.

**FIG. 4.**
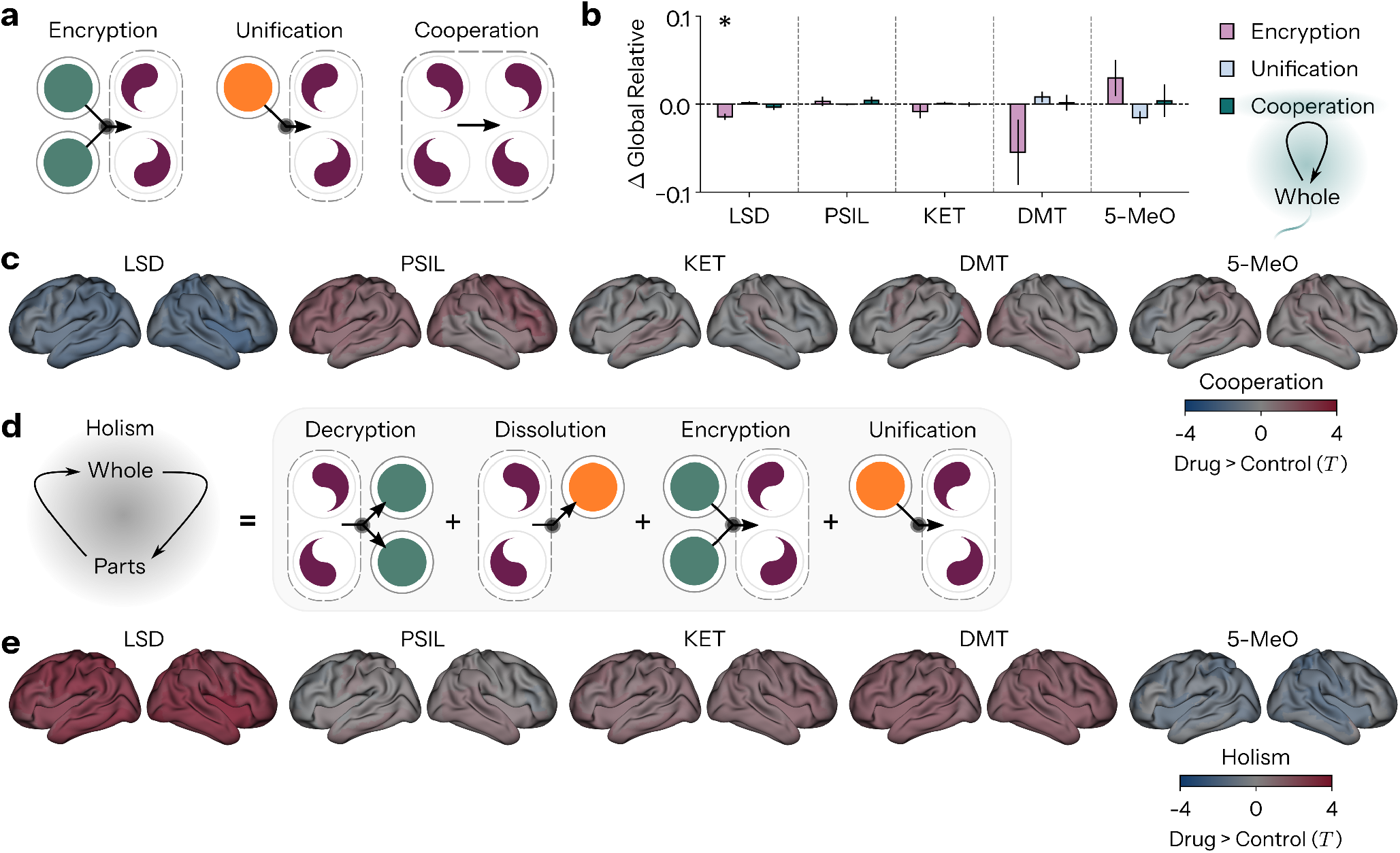
Effects of psychedelics on higher-order relative information dynamics. **a** Information dynamics invisible to existing measures: Encryption, Unification, Cooperation. **b** Global relative changes in Encryption, Unification, Cooperation (10-plets). **c** Topographic statistical maps for regional differences in relative Cooperation (10-plets). **d** A novel aggregate phenomena: Holism. Holism measures how much information is moved between the parts and the whole, thus quantifying the total traffic of information between scales, the total transformation of information between lower-order and higher-order formats. Holism measures the sum of Decryption, Dissolution, Encryption, and Unification. **e** Topographic statistical maps for regional differences in relative Holism (10-plets). (* indicates *p <* .05, FDR-corrected Benjamini-Hochberg procedure. Bars indicate mean *±* SEM.)

Hence, we continue our analysis by examining relative alterations in these previously undetected information dynamics, with absolute changes presented in Supplementary Fig. S3. Moreover, since existing measures were confined to the space of lower-order (univariate or bivariate) interactions, and our framework is not, we showcase higherorder global results, constructing multivariate sources and targets with each element comprising five brain regions (rather than one). We provide relative global changes for 2-plets in Supplementary Fig. S3. We find that global relative Encryption is decreased under LSD (*p*_FDR_ = 0.001; Fig. 4b), with no significant effects for psilocybin, ketamine, DMT, or 5-MeO-DMT. Unification does not reach significance for any drug (LSD *p*_FDR_ = 0.058, PSIL *p*_FDR_ = 0.915, KET *p*_FDR_ = 0.403, DMT *p*_FDR_ = 0.251, 5-MeO-DMT *p*_FDR_ = 0.078). Cooperation likewise did not reach significance for any psychedelic, globally or regionally (Fig. 4b,c; LSD: *p*_FDR_ = 0.597; PSIL: *p*_FDR_ = 0.597; KET: *p*_FDR_ = 0.948; DMT: *p*_FDR_ = 0.948; 5-MeO-DMT: *p*_FDR_ = 0.948). Full regional results for absolute and relative values can be seen in Supplementary Figs. S5 and S7.

Given our comprehensive access to the information atoms that compose time-evolving interactions in the brain, we can now flexibly recombine them to construct, from the bottom up, new measures for addressing specific questions. Contemporary neuroscience increasingly emphasizes linking levels of organisation, and reconciling scale-specific notions of brain function [44, 45]. One avenue in this research program that has been overlooked however, is the formal and empirical understanding of when, and to what extent, scales are integrated in the brain [46]. More concretely, there is an absence of measures that specifically quantify the total information flow between the parts and the whole of brain systems. Thus, to probe the question of whether psychedelics induce scale integration in the human brain, we introduce a measure of the amount of information being transferred between scales, termed *Holism*, defined as the sum of Decryption, Dissolution, Encryption, and Unification (Fig. 4d). Fascinatingly, we find that no compound significantly increased global relative Holism (LSD: *p*_FDR_ = 0.056, PSIL: *p*_FDR_ = 0.749, KET: *p*_FDR_ = 0.437, DMT: *p*_FDR_ = 0.388, 5-MeO-DMT: *p*_FDR_ = 0.437). Topographically however, LSD shows a robust, near-global regional increase (Fig. 4e) (76 of 78 regions significant after TFCE correction, all positive), peaking in cuneus (BF_10_ = 22.1), paracentral lobule (BF_10_ = 14.1), and Rolandic operculum (BF_10_ = 12.6). Despite this widespread regional effect, no other drug shows any regionally significant change in Holism.

Therefore, we find that psychedelics do not, across the board, robustly increase the relative prevalence of emergent higher-order information integration in large-scale human brain activity.

### D. Converging spatial patterns of psychedelic effects

Given the common and distinct effects of psychedelics on information dynamics, we next examine the topo-graphic organisation of this restructuring across drugs.

We performed a random-effects meta-analysis for all ten distinct types of regional relative information dynamics (see Methods), obtaining a pooled *d* value per cortical parcel per drug (Fig. 5a, inflated surface). Additionally, we provide discrete results for the number of modes of information dynamics that possess *p <* .05 for each region per drug (Fig. 5a, flattened surface). Examining the regions most affected by each drug, we find that for LSD the strongest effects are inferior occipital (*d* = 0.637), for psilocybin the inferior frontal gyrus (*d* = 1.044), and for ketamine the mid-cingulum (*d* = 0.649). For DMT, the strongest effects reside in the right superior parietal gyrus (*d* = 0.973), and for 5-MeO-DMT in the right middle occipital gyrus (*d* = 0.984). While these hotspots are distinct across drugs, the parahippocampus, and right calcarine, lingual, and superior occipital gyri each show at least four significant modes of information dynamics in three or more drugs. This occipital commonality is driven predominantly by the serotonergic psychedelics (LSD, psilocybin, DMT, 5-MeO-DMT), with ketamine contributing only via parahippocampal cortex.

**FIG. 5.**
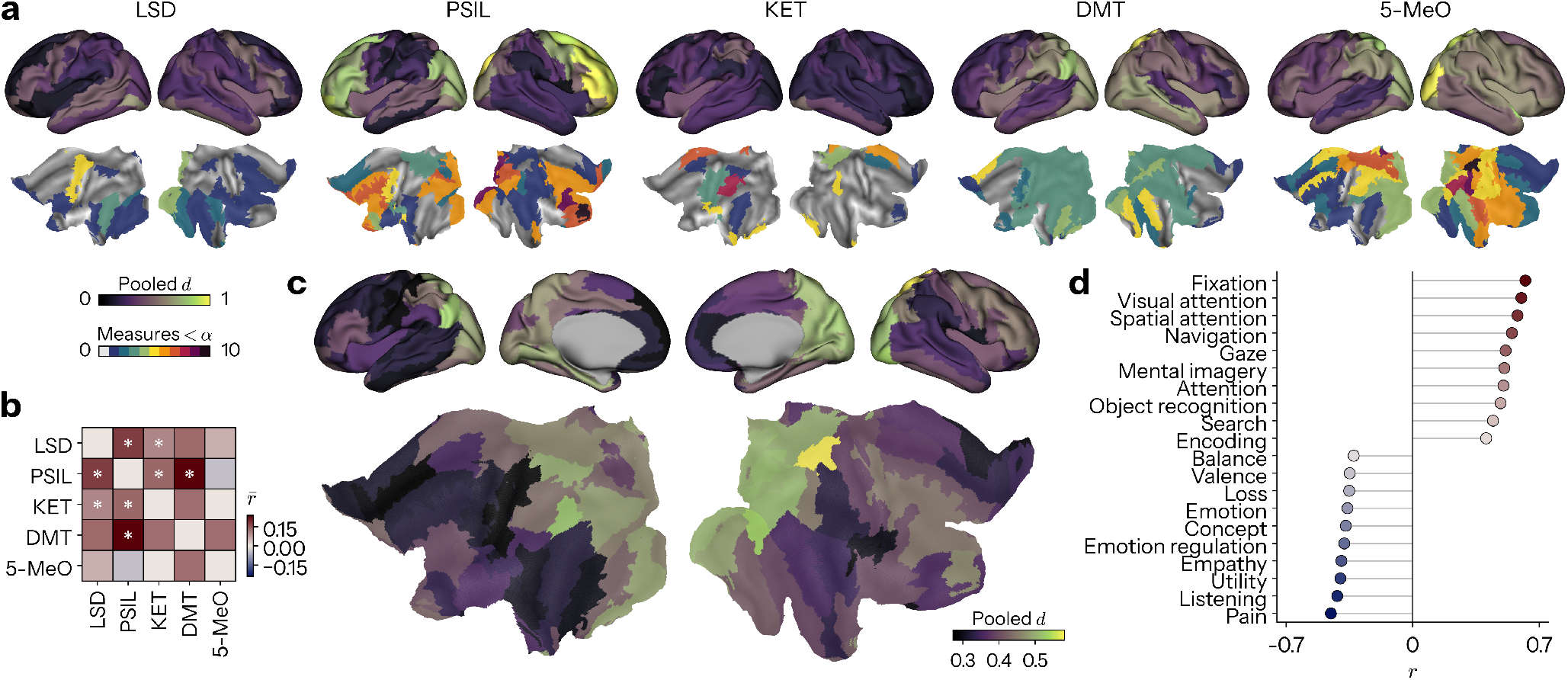
Topographic organisation of psychedelic modulation of information dynamics. **a** Cortical distribution of psychedelic effects pooled across all modes of relative information dynamics. First row: inflated cortical surfaces depict random effects meta-analytic pooled effect size. Second row: flattened cortical surfaces depict the number of significant (*p <* .05) alterations in information dynamics. **b** Mean correlation of statistical contrast maps across drugs. **c** Cortical distribution of psychedelic effects pooled across all drugs and modes of information dynamics. **d** Correlation of fully pooled meta-analytic map from **c** with meta-analytic cognitive maps from Neurosynth. The top ten most correlated and anti-correlated terms are displayed. (* indicates *p <* .05)

To take into account the direction, rather than just the magnitude of effect, and examine how the topographies of alterations in information dynamics align between drugs, we correlate the *T* -statistic maps for each mode of information dynamics across drugs (Fig. 5b). The strongest associations on average were expectedly shown between psilocybin and DMT 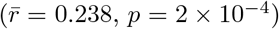, as well as between LSD and psilocybin 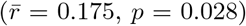. Consistent with its radically singular subjective experience, 5-MeO-DMT shows no gross significant association with any other drug.

When performing the meta-analysis pooling across all drugs and modes of information, we find that the cortical area most affected by psychedelic drugs is the right superior parietal gyrus (*d* = 0.581; Fig. 5c). To glean context for these resting-state alterations, we assay via reverse inference their correspondence to 123 cortical cognitive term maps obtained via Neurosynth (see Methods), a meta-analytic database of more than 14,000 neuroimaging experiments [47] (Fig. 5d). Full list of terms can be found in Supplementary Table S1. Interestingly, the most strongly positively associated terms broadly reflect visuospatial and executive processing (Top 3; fixation: *r* = 0.62, visual attention: *r* = 0.60, spatial attention: *r* = 0.58), while the most strongly negatively associated terms reflect no-ciceptive and social processing (Top 3; pain: *r* = *−*0.45, listening: *r* = *−*0.42, utility: *r* = *−*0.40). Results for each drug separately are shown in Supplementary Fig. S8.

### E. A minimal generative model of psychedelic effects

Finally, we turn our attention to the mechanistic basis of these effects. That is, the causal relations that produce the informational profile of the psychedelic brain. To do so, we employ the minimal setup of a two-node vector autoregressive (VAR) process subject to Gaussian noise (Fig. 6a). The model includes three main parameters: self-coupling (*a*), which determines the influence of a node over itself, cross-coupling (*b*), which determines the influence of a node on the other node, and noise correlation (*c*), which determines the similarity of the externally inputted noise to each node (see Methods).

**FIG. 6.**
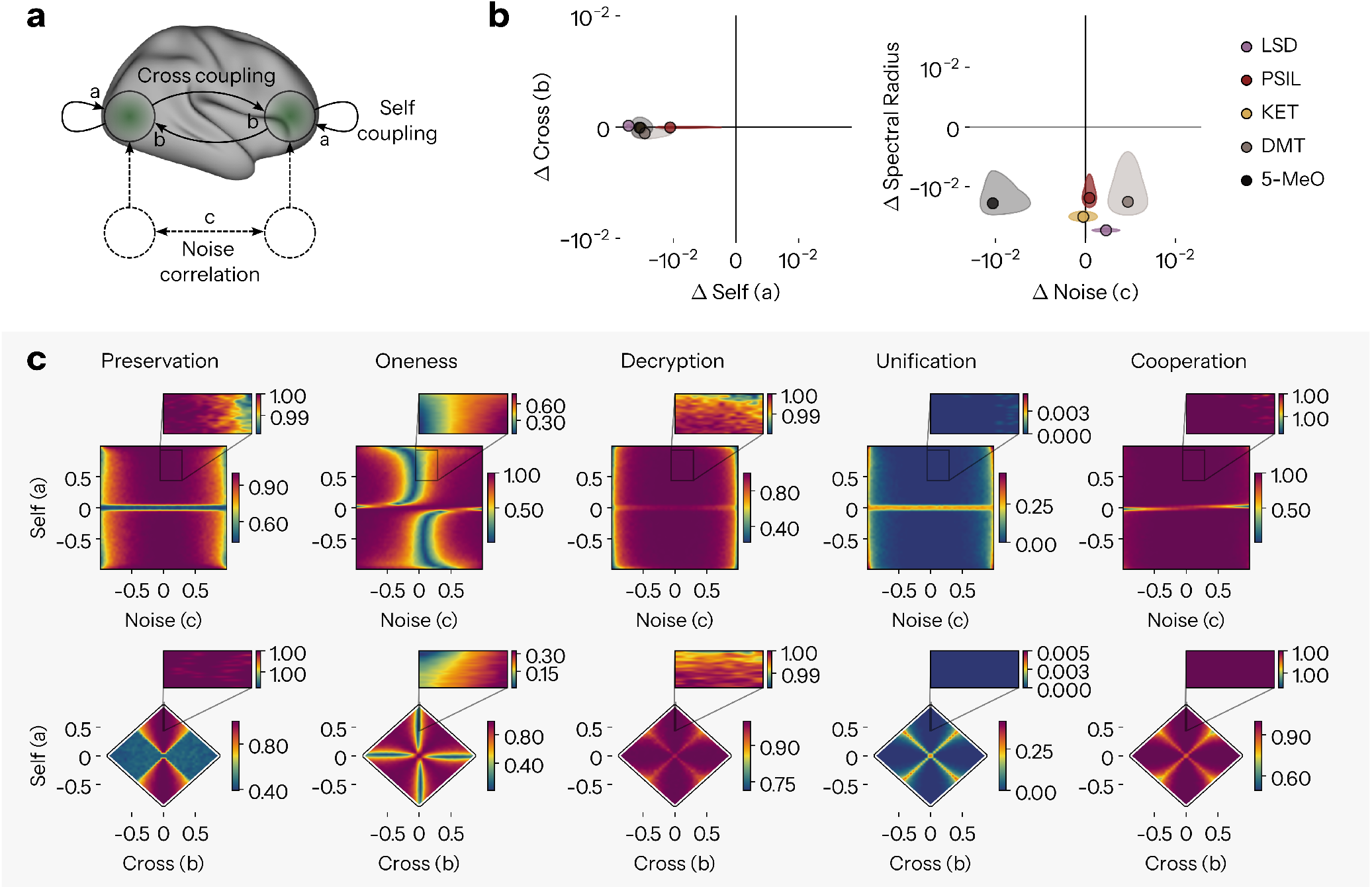
A minimal generative model of psychedelic effects. **a** Two node deterministic vector autoregressive (VAR) process subject to Gaussian noise possessing three parameters: Self-coupling (*a*), which determines the influence of a node over itself, Cross-coupling (*b*), which determines the influence of a node on the other node, and Noise (*c*), which determines the correlation of the externally inputted noise to each node, in other words the shared innovation between neural populations. **b** Average changes in parameters of the model fitted to empirical data from control to drug condition. Left plot: self-versus cross-coupling (*a* versus *b*). Right plot: noise correlation versus the derived spectral radius of the coupling matrix (*c* versus *ρ*). Dots represent mean, shaded ellipses represent SEM across participants. **c** Reconfigurations in relative information dynamics in the sweep of model parameters. First row: *a* versus *c* with fixed *b* = 0.02. Second row: *a* versus *b* with fixed *c* = 0.06. ΦID is computed numerically from the full covariance matrices of the system. *a* vs *b* plot has black borders denoting the boundary of model parameters that yield valid covariance matrices. Zoomed inlets display the space occupied by mean empirical coefficients.

With these models fitted to our empirical data (see Methods), we can examine the differences in coefficients between conditions (Fig. 6b). Taking global averages, we find that psychedelics ablate self-coupling, with the strongest effects for LSD and ketamine (LSD: BF_10_ = 3.01*×* 10^6^, PSIL: BF_10_ = 0.498, KET: BF_10_ = 110.4, DMT: BF_10_ = 0.641, 5-MeO-DMT: BF_10_ = 1.99). On the other hand, cross-coupling remains relatively unchanged across all five drugs (LSD: BF_10_ = 0.381, PSIL: BF_10_ = 0.298, KET: BF_10_ = 0.308, DMT: BF_10_ = 0.434, 5-MeO-DMT: BF_10_ = 0.238). Examining the spectral radius (*ρ*) of the coupling matrix, an index of the temporal stability of the system, we find that drugs tend to exhibit lower *ρ*, particularly strongly so for LSD and ketamine (LSD: BF_10_ = 4.42 *×*10^6^, PSIL: BF_10_ = 1.087, KET: BF_10_ = 117.0, DMT: BF_10_ = 0.519, 5-MeO-DMT: BF_10_ = 0.632). Finally, noise correlation shows no consistent direction across drugs: it trends upward for LSD and DMT, but interestingly downward for ketamine and 5-MeO-DMT (LSD: BF_10_ = 0.599, PSIL: BF_10_ = 0.304, KET: BF_10_ = 0.241, DMT: BF_10_ = 1.339, 5-MeO-DMT: BF_10_ = 1.758). Full statistics can be found in Supplementary Table S2. Thus, we find that, on average, psychedelics reduce self-coupling, most robustly for LSD and ketamine.

To confirm and explore these effects further, we perform a sweep of model parameters (Fig. 6c). Following the same logic as the empirical scenario, it is trivial that the model will produce results that display different levels of information dynamics purely driven by changes in the TDMI. Accordingly, we investigate relative (normalised) results here (Fig. 6c), providing absolute (unnormalised) results in Supplementary Fig. S9. We display sweep results here for five modes of information dynamics – Preservation, Oneness, Decryption, Unification, and Cooperation – since these cover the ground of driving factors for existing measures as well as including novel measures of integration. Results for all modes of information dynamics can be found in Supplementary Fig. S10.

For Preservation, the model separates two effects. The loss of absolute Preservation, which drives the rise in entropy rate, is explained almost entirely by the fall in self-coupling *a*: the less a region constrains its own future, the less information it keeps to itself. Relative Preservation, by contrast, is largely insensitive to *a* and instead depends on contemporaneous linkage between regions *c*, so input correlation reduces the share of information that regions store privately, even when the total amount is unchanged. Loss of Encryption, on the other hand, is tied to reduced *a* in both absolute and relative terms. We find a heterogeneous sensitivity in Oneness, which can increase as a consequence of any one of changes of *a, b* or *c*. In other words, changes in FC can have multiple concurrent or isolated mechanisms, and may be driven by increases in cross-coupling, or on the other hand, reductions in self-coupling or increases in noise. This heterogeneity also offers a candidate account of 5-MeO-DMT’s distinctive reversal of this pattern: 5-MeO-DMT shows a reduction rather than an increase in noise correlation, and since Oneness rises steeply with *c* in the model (with Preservation falling weakly), this offers a plausible mechanism for its uniquely significant decreased Oneness and increased Preservation. In sum, simple changes in self-coupling and shared input can underlie alterations in complex information dynamics, without any change in the cross-coupling between regions.

## III. DISCUSSION

Our investigation establishes the informational principles that govern psychedelic-induced alterations in large-scale human brain dynamics. Specifically, we provide a framework to detect and interpret the information atoms composing functional interactions in the brain, and employ this to reveal the overlapping basis functions that drive prior empirical results. We demonstrated how psychedelics affect multiple, until now, unexplored forms of information dynamics, and that overall, while the effects of psychedelics are complex and heterogeneous, they can be accounted for by a minimal set of mechanisms.

We showed that increases in complexity via entropy rate can be considered instead as a decrease in active information storage (AIS). While this may seem mathe-matically trivial, making it explicit empirically encourages different intuitions for theory generation. For example, one popular hypothesis posits an identity in which gains in entropy rate are a corollary to the felt ‘richness’ [13] or ‘expansiveness’ [48] of a mental state. However, if one were to invest the same interpretative weight in the concomitant loss of stored information, would it not be expected to engender something counter to this effect? One could argue this interruption of informational continuity may corrupt a “stream” of consciousness, preventing representational fidelity, mnemonic inference, and semantic depth. What’s more, we revealed that the AIS is in fact the sum of Erasure, Copy, Oneness and Preservation, four totally qualitatively distinct modes of information dynamics. In this way, it would be prudent to consider how states with the same change in entropy rate, but totally different underlying information dynamics being ablated, may differ.

We revealed that Oneness is the information dynamical foundation of functional connectivity (FC). This shows the dominant paradigm for inferring inter-regional signaling in neuroscience reflects only one of many possible modes of information dynamics [22, 49]. In light of this, we may interrogate the construct validity of FC as originally conceived – as a tool to distinguish neural assemblies [50]. While Oneness formally captures key hypothesised properties of neural assemblies, such as short-term memory and resilience to individual element failures, it is insensitive to the critical notion that assemblies possess emergent properties at the collective level that are absent in their individual parts [50]. Though the identification of FC as Oneness does not reduce the distance between association and causation in FC research, it provides clarity that can ground the interpretation of FC results beyond the ambiguous reference to its indication of “statistical dependency” [51–53]. This discussion is particularly pertinent to the present context, since a sizable portion of theory generation and testing in the neuroscience of psychedelics occurs through the use of FC. Thus, regardless of how elaborate the algorithms applied to resultant FC matrices, they are still fundamentally built from Oneness, meaning prior claims regarding functional network desegregation and whole-brain integration can only be made for this single form of information dynamics [5, 8].

Examining information transfer via directed functional connectivity (dFC), we show that classic measures (TE) capture the sum of Copy, Migration, Decryption, and Dissolution. While the Copy and Migration align with conventional interpretations of dFC, the latter two may appear more intriguing, since rather than measuring the exchange of information between regions per se, they measure the transfer of information from the whole to the parts (from macro to micro). The recognition of these forms of information dynamics in their own right is imperative, since in numerical terms they are the ones most strongly globally altered across these drugs, but, since they change in opposing directions, their effects are canceled out in dFC. For their complementary processes, Encryption and Unification, we saw again this push-pull effect. Our results therefore hint at the occurrence of a specific computational scheme under psychedelics, whereby distributed de-localized representations are dynamically recast out of and into independent representations, rather than undifferentiated multi-localized representations.

Our results revealed the confluence inherent to methods used in contemporary neuroimaging, particularly in the neuroscience of psychedelics. That is, we showed how the informational basis of FC is a component of the information storage that drags down entropy rate, and further that information storage and transfer, despite trying to measure distinct phenomena, overlap in measuring the duplication of information via Copy. But beyond this methodological point, our results open up a striking dilemma regarding how we conceptually organize our empirical understanding of brain function, and how psychedelic drugs alter it. Specifically, it is clear that the parlance of “integration” or “complexity” on their own will not suffice, as almost any one of the modes of information dynamics presented herein could, if desired, be described as a manifestation of either of these phenomena. For instance, claims that psychedelics increase “integration” in the brain require careful qualification: while this may be true for relative Oneness (with the exception of 5-MeO-DMT), it is not so evident for something like Co-operation. Similarly, “complexity”, often simply construed as a kind of dynamical diversity, in which the future looks unlike the past, could theoretically incorporate any temporally asymmetric form of information dynamics, thus only excluding Preservation, Oneness and Cooperation. Ironically, in complexity science more broadly, when the concept of “complexity” is invoked, it is precisely referring to the kinds of hierarchical emergence that Cooperation is suggestive of [54, 55]. Complicating matters further, our computational model showed that changes in information integration across regions need not stem from a shift in the coupling between those regions, but can instead arise from changes in how each region couples to itself.

With this in mind, a different story for the large-scale brain action of psychedelics begins to emerge. Loss of absolute Preservation indicates that psychedelics disrupt the ability of cortical populations to locally encode information. Private self-dependence is ablated, and unique representations fail to consolidate. Instead, in a deviation from independence, increases in relative Oneness tell us that the brain spends its informational budget ensuring content is available from anywhere, all the time. In this way, psychedelics can engender a regime of persistent “hyper-representation” in the brain. Intriguingly however, 5-MeO-DMT causes precisely the opposite of these two effects, by bolstering Preservation and diminishing Oneness, it elicits a unique state of local stability and global hypo-representation. Taken together, psychedelics engender a situation in the human brain whereby information can be made everywhere, depending on nowhere. Wherein the balance between multi-localised and delocalised formats can also shift, even without an overall increase in the traffic across space and scales. Put concisely, the psychedelic brain is one in which cortical populations can become alike, but not necessarily together.

Consistent with a very recent analysis from the ‘BOLD psychedelic consortium’, of eleven acute rsfMRI datasets covering four psychedelic drugs [5], we observe drastic heterogeneity in the neural effects of different compounds. However, like this study, we are unable to discern whether this reflects genuine distinctions between drugs, contesting monolithic framings of ‘*the* psychedelic brain’, or instead reflects differences in study design and data collection. For example, while the LSD, psilocybin and ketamine data are MEG, the DMT and 5-MeO-DMT data are EEG. Thus, despite our data-harmonization, differential in-plane and depth resolution, sensitivity to tangential versus radial currents, and conductivity are present. Additionally, all drugs except 5-MeO-DMT employ a saline placebo group. Then again, the inevitable unblinding when ingesting a powerful psychoactive drug has largely led to the conclusion that such placebo *groups* are likely not placebo *controls* properly construed [56–58]. Another contributing factor here, is likely that we are analysing so-called ‘resting-state’ data, which has been argued to be an unconstrained and artificial construct, that is not representative of a native physiological or psychological state, nor engaging of the breadth of cognitive processes that occur in typical complex environments, and thus acts as an unreliable proxy for latent functional organisation [59–62]. Furthermore, given that different psychometric tools were used to assess the nature of participants’ experiences in each study, we are unable to use this data in a robust way to wrangle potentially incomparable cognitive operations, and explicitly link neural and subjective features across drugs. More broadly, our analysis approach suffers from a number of limitations. At the outset, our computational setup involves assuming that brain dynamics are Gaussian and Markovian. Importantly, given that our decomposition is fundamentally an under-determined algebraic framework with no unique solution, demanding the definition of a double redundancy function, it is unclear the extent to which the proposition of an alternative definition may alter results. Further, we also restrict our analysis to the case of two variables at a time, since the decomposition lattice grows with the square of the Dedekind number of elements (*D*(*k*) *−*2)^2^, meaning a nine element system would produce more information atoms than physical atoms in the observable universe (*>* 10^82^).

Our work opens the door for multiple avenues for future work. First, it will be important to investigate how these electrophysiological results translate to other non-invasive neuroimaging techniques. Specifically, given ongoing controversy around psychedelics’ ability to dissociate hemodynamic responses from neural activity [63–65], it is unclear whether our results will replicate in fMRI. Second, it will be important to understand how local and global changes in information dynamics correspond to the post-acute psychological effects of psychedelics. This also raises the possibility that therapeutic interventions, like drugs, could be designed to target particular modes of information dynamics. More broadly, it is transparent that our approach can be applied far beyond the confines of the neuroscience of psychedelics. For instance, one could ask what drives the loss of signal diversity that marks loss of consciousness during sleep, anaesthesia, or coma [66], given it must involve Erasure, Copy, One-ness, or Preservation. One could investigate whether in general working memory is subserved more by Preservation (unique information storage), Oneness (redundant information storage), or Cooperation (synergistic information storage) in cortico-hippocampal interactions, and whether this differs depending on whether the memory is declarative or not [67, 68]. Can different hierarchical coding schemes in primary sensory cortices reconcile their views on the nature of feedback signals if thinking in terms of information rather than amplitude? whereby both suppression and amplification reflect a common form of dependency? [69] These examples illustrate that the methodology and vocabulary presented herein offer a significant opportunity for the theoretical refinement and formal testing of central questions in neuroscience [21].

In conclusion, our results challenge the coarse view that psychedelics increase “global integration” in the human brain. Instead, we build the basis for a more involved perspective, where the private self-dependence of cortical populations is generally thwarted, but truly collective interactions are not enhanced. Cortical populations can indeed persistently align, but by mirroring or matching each other, not by acting with persistent complementarity.

## IV. METHODS

### Data

We used previously obtained M/EEG data from five classical and atypical psychedelic compounds: lysergic acid diethylamide (LSD) [70], 4-phosphoryloxy-N,N-dimethyltryptamine (psilocybin) [71], ketamine [72], N,N-dimethyltryptamine (DMT) [27], and 5-methoxy-N,N-dimethyltryptamine (5-MeO-DMT) [42]. Full experimental, acquisition, and preprocessing details are provided as Supplementary Material (Supplementary Section S1), but are summarized here in brief. These are within-subject resting-state M/EEG recordings in healthy participants, each comprising a sober control condition and a peak subjective effect drug condition. Data were harmonized by source localizing and parcellating to the same atlas (AAL-90) [73] with subcortical regions removed; then bandpass filtering to 1–50 Hz with a zero-phase 2nd-order elliptic filter, downsampling to 100 Hz with integer decimation, truncating epochs to the minimum length of 2 s, and demeaning. Epochs were treated as stationary realizations of an underlying stochastic process in all analyses.

### Integrated Information Decomposition

Integrated Information Decomposition (ΦID) is a mathematical framework for evaluating how information is structured, and how this evolves over time, in complex dynamical systems [20]. ΦID advances upon previous information decompositions by differentiating the role of each source and each target when partitioning the total amount of information flowing from past to future. In other words, ΦID comprehensively stratifies the structure of cause (forward) and effect (backward) information simultaneously, yielding a complete taxonomy of how information is stored, transferred, and transformed over time.

Treating the past activity of a pair of brain region assources 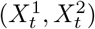, and the future of those same regions as targets 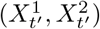, we use ΦID to decompose their TDMI 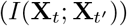 into information “atoms” 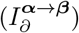:

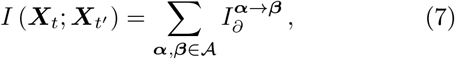

where *A* indicates the set of possible collections of sets of sources. For instance, in the case of two variables *X*^1^ and *X*^2^, the set *A* is given by

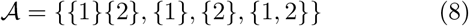

which represent, respectively, the redundant information shared by *X*^1^ and *X*^2^, the information uniquely provided by *X*^1^, the information uniquely provided by *X*^2^, and the synergistic information that can only be obtained by jointly considering both variables. Hence, the atoms 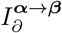 represent any symmetric or asymmetric combination of redundant, unique or synergistic information in the past (***α***) and future (***β***), forming the double decomposition *→* ***α****→* ***β****∈ A × A* . For example, 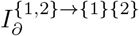 refers to synergistic information that is transformed into redundant information over time, and 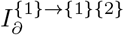 refers to unique information in the first variable being transformed into redundant information over time. We refer to these atoms as *S →R* and *U*_1_ *→R* for simplicity.

To perform the decomposition, ΦID introduces the so-called “double redundancy lattice”, i.e. the partially ordered set defined by the cartesian product *A × A*. In practice, this provides a system of fifteen linear equations that comprise standard Shannon mutual information and intersection information expressed in terms of information atoms. This lattice structure entails that one can solve the system and derive values for each atom through the Möbius inversion:

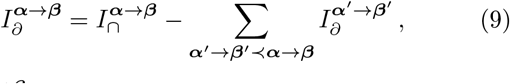

where 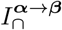 is the intersection information, which reduces to mutual information in the case of a single source. More literally, taking *S →R* as an example again, ΦID allows us to say:

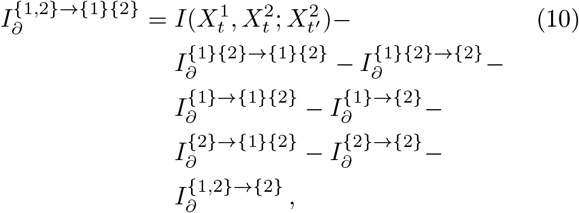

and similarly for *U*_1_ *→ R*:

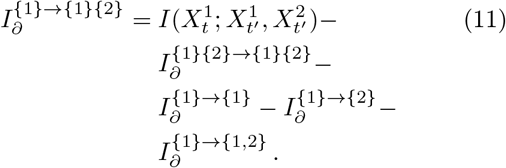

Hence, this enables one to decompose existing mutual information-based quantities, such as Transfer Entropy and Active Information Storage, simply by explicitly writing their informational contributions via Eq. (9). For AIS, we can therefore state:

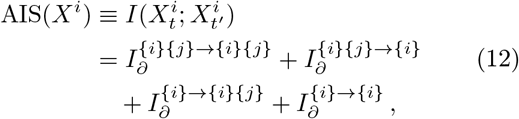

whereas TE reads:

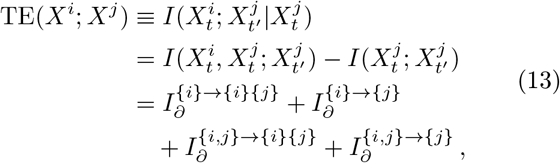

where *j* denotes the other variable in the ΦID decomposition. These decompositions correspond to that of Eqs. (2)-(6).

As just hinted however, ΦID is a system of fifteen equations aiming to resolve sixteen atoms, we must first define a starting point 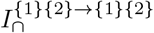, or double-redundancy function, for which^*∩*^we use a direct extension of the minimum mutual information (MMI) [20, 74]:

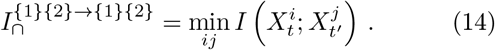

With these definitions at hand, we first compute ΦID on pairs of brain regions (2-plets), then account for effects at greater scales by computing ΦID on multivariate sets five regions split in a bipartite scheme (10-plets).

### VAR models

To effectively model the evolving inter-dependencies of neural dynamics, we employ vector autoregressive (VAR) models. VARs are sets of linear equations capturing the deterministic growth of activity states subject to noise:

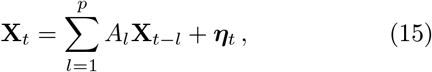

with shorthand matrix notation:

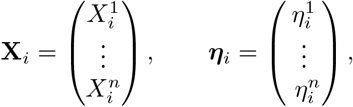

where *p* is a positive integer representing the model order, or the length of incorporated history, ***η*** is a multivariate white noise term, and *A*_*l*_ is the *n× n* evolution matrix containing the *l*-th lag coupling coefficients.

Fitting these models to pairs of brain regions in our empirical data, we follow standard practice by estimating *A*_*l*_ using ordinary least squares via QR decomposition, and extracting the noise covariance matrix *V* as

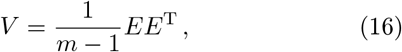

where *E* is the matrix of the residuals, and *m* the number of samples (*T− p*).

Put simply, *V* thus captures the instantaneous coupling, shared innovation, or noise correlation, between neural populations. This is in contrast to the lagged causal interactions of *A*_*l*_, the diagonal entries of which determine lagged self-coupling, and the off-diagonals which determine the cross-coupling.

To study information dynamics in VAR models via the covariances of the constituent time-lagged variables, we must first gain access to the elements of the autocovariance matrices 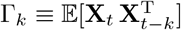. To do so, we rewrite Eq. (15) as:

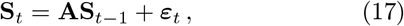

with matrix notation for the elements:

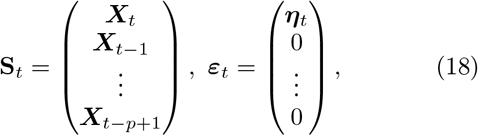

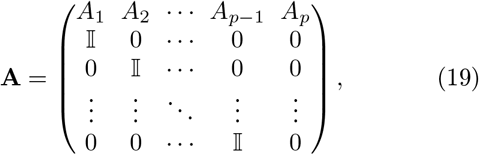

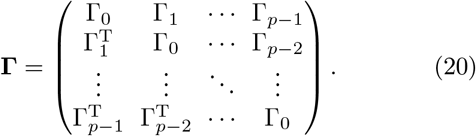

From here, it follows that **Γ** must satisfy the discrete Lyapunov equation [75]:

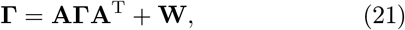

where **W** is a block matrix with residual covariance V in the first entry and zeros elsewhere.

Hence, once *A*_*k*_ and *V* are known, it is possible to obtain the autocovariance matrices from Eq. (21) and thus the full covariance of the system for gaussian information-theoretic treatment. For example, the TDMI in the context of a VAR model is simply:

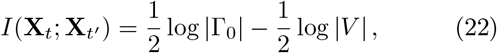

with other mutual information terms being recovered analogously by marginalizing particular elements in the covariances.

In our mechanistic simulations, we explore the parameter space of valid covariances of a symmetric modular reduced VAR(1) system of the form:

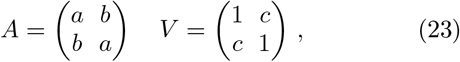

where *a* is the self-coupling, *b* is the cross coupling, and *c* is the noise-correlation. We perform a parameter sweep over these values across a grid *S*_*a*_*× S*_*b*_*× S*_*c*_ given by intervals *S*_*a*_ *∈* [*−*0.99, 0.99] (*N* = 50), *S*_*b*_ *∈* [*−*0.99, 0.99] (*N* = 50), and *S*_*c*_ *∈* [*−*0.99, 0.99] (*N* = 50), performing ΦID in each scenario, and normalizing quantities, just like our empirical values, as described in the next section.

### Null models

The promise and peril of directly comparing information-theoretic measures across different systems and scenarios lies in their intrinsic sensitivity. Even systems with similar structural or statistical properties can vary widely in their observed joint, conditional, and marginal entropies, making direct head-to-head comparisons challenging. This issue is particularly pronounced in information decomposition frameworks, where the values of information atoms are inherently constrained or inflated by the quantity from which they are decomposed [41]. Thus, in order to rigorously compare these quantities, a normalization procedure is required.

To this end, we develop a null model procedure for ΦID, extending previous work, by constructing null distributions from systems with equivalent TDMI [41]. Specifically, we construct VAR models with random evolution matrices *A*, and then globally rescale them to control for their spectral radius *ρ*, which determines the TDMI. To ensure that the null distributions remain meaningfully close to those of the real systems, the matrices *A* are sampled from the set of empirical evolution matrices obtained by fitting VAR models to the control condition data, denoted as *P* (*A*). In other words, we create a suite of models with quasi-arbitrary causal structures, but with their overall stability, and thus information flowing from past to future, being explicitly set. Thus, in practice, once the TDMI of the empirical system *p*(*X*) is known, for each null model *q*_*i*_(*X*) we sample *A*_*k*_*∼ P* (*A*) and 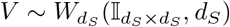, with *d*_*S*_ being the size of the system of Eq. (17), and then optimize *ρ* through Eqs. (21)-(22) to satisfy our constraint on the TDMI. For each pair of brain regions, we generate 1000 null systems, perform ΦID on them, and normalize both individual atoms and the aggregates of atoms by taking their quantiles w.r.t. the null distribution. This procedure ensures that any obtained difference reflects a genuine structural or dynamical distinction, rather than trivial variations in mutual information, thus conveying the relative contributions of each information atom to the overall information present in the system [41].

### Spatial statistics

To correct for multiple comparisons across the 78 cortical regions in each regional contrast, we used threshold-free cluster enhancement (TFCE) [76], applied via signflip permutation over the cortical adjacency structure of the AAL-90 atlas. For each region *v* and permutation, the TFCE score is Σ_*h*_ *e*(*v, h*)^*E*^ *·h*^*H*^ *·δh*, where *e*(*v, h*) is the extent of the supra-threshold cluster containing *v* at threshold *h, δh* = 0.2 is the threshold step, and *H* = 2, *E* = 2*/D*, where for our cortical surface the intrinsic dimensionality *D* = 2. Null TFCE scores were generated from 5000 sign-flip permutations, and corrected *p*-values (*p*_TFCE_) were obtained region-wise from the resulting null distribution, family-wise across all regions.

### Meta-analysis

To examine the spatial layout and overlap of psychedelic effects on cortical information dynamics, we performed a random-effects meta-analysis using the DerSimonian-Laird method. Specifically, for each drug-region combination, we pooled Cohen’s *d* effect sizes across the 10 information measures using inverse-variance weights:

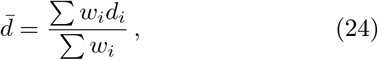

where *d*_*i*_ is the effect size for the *i*-th mode of information dynamics. Within-measure variance was estimated as 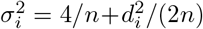, where *n* is the number of participants, and thus fixed-effect weights 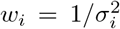 were used to compute an initial 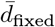. Between-measure heterogeneity was then estimated as:

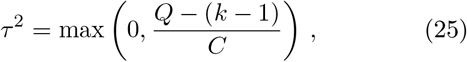

where 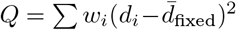, *k* is the number of measures, and 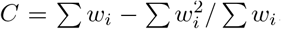. Thus, the final random effects pooled estimate used weights 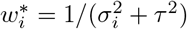 in Eq. (24), can be derived with standard error 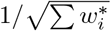. To identify regions showing consistent effects across compounds, we repeat this procedure pooling all drug-measure combinations within each region.

To contextualize our meta-analytic maps, we test their association with meta-analytic maps from Neurosynth [47]. The volumetric *P* (function |activation) association test maps for the 123 terms that overlap between Neurosynth and the Cognitive Atlas [77] were used, parcellated into the same atlas as our empirical data.

## V. DATA AVAILABILITY

This study analyses previously published human M/EEG datasets [27, 42, 70–72]. Data are available upon reasonable request from the corresponding authors of the primary studies, subject to their respective original data-sharing policies and permissions.

## VI. CODE AVAILABILITY

Code is publicly available at github.com/ gblackburne/infodelia.

## VII. ACKNOWLEDGEMENTS

G.B. is supported by the Leverhulme Trust. J.I.S. is supported by Wellcome Leap. P.M. and P.A.M. are supported by Tiny Blue Dot Foundation.

## VIII. CONTRIBUTIONS

G.B.: Conceptualization, Methodology, Software, Formal analysis, Investigation, Data curation, Writing - original draft, Writing - review & editing, Project administration. A.L.: Conceptualization, Methodology, Software, Formal analysis, Writing - review & editing. P.M.: Writing - review & editing. R.G.M.: Writing - review & editing. R.L.C.-H.: Writing - review & editing. C.T.: Writing - review & editing. S.M.: Writing - review & editing. J.I.S.: Supervision, Writing - review & editing. P.A.M.M.: Supervision, Writing - review & editing

## IX. ETHICS DECLARATIONS

Each of the five contributing studies were approved by the relevant research ethics authority and conducted in accordance with the Declaration of Helsinki (REC references: LSD, 13/LO/0229; psilocybin, 08/H0101/144; ketamine, 13/WA/0059; DMT, 16/LO/0897; 5-MeO-DMT, 19437/004). All participants provided written informed consent. Full ethical statements, details of approval, and consent procedures are provided in Supplementary Section S1 and the respective original publications [27, 42, 70–72].

## X. CITATION DIVERSITY

Recent work in several fields of science has identified a bias in citation practices such that papers from marginalised demographics are under-cited relative to the number of such papers in the field [78–80]. To report transparently on our own bias, first, we classified the gender of the first names of the first and last authors [78]. Our references contain 0.5% woman(first)/woman(last), 6.94% man/woman, 15.57% woman/man, and 76.99% man/man. Second, we performed a similar procedure for obtaining predicted racial or ethnic category. By this measure, splitting into inferred white authors (W) and authors of colour (C), our references contain 7.55% C/C 13.66% W/C, 19.22% C/W, and 51.61% W/W. We note that these algorithmic proxies are limited, cannot infer the full spectrum of self-determined gender identity, structurally excludes non-binary, transgender, and intersex researchers, and cannot adequately account for Indigenous and mixed-race authors. We look forward to future work that could help us to better understand how to support equitable practices in science [81].

