## Supplementary material for "Psychedelics restructure information in the human brain": Suplementary Material

### Supplementary Material
### Psychedelics restructure information in the human brain

George Blackburne,[1,2,*] Alberto Liardi,[2] Pablo Mallaroni,[2,3] Rosalind G. McAlpine,[4,5] Robin L. Carhart-Harris,[6,7] Christopher Timmermann,[1,7] Suresh Muthukumaraswamy,[8] Jeremy I. Skipper,[1] and Pedro A. M. Mediano[2,1]

[1] *Department of Experimental Psychology, University College London*
[2] *Department of Computing, Imperial College London*
[3] *Department of Neuropsychology and Psychopharmacology, Maastricht University*
[4] *Clinical Psychopharmacology Unit, University College London*
[5] *Department of Clinical, Educational, and Health Psychology, University College London*
[6] *Department of Neurology, University of California San Francisco*
[7] *Centre for Psychedelic Research, Department of Brain Sciences, Imperial College London*
[8] *Faculty of Medical and Health Sciences, School of Pharmacy, University of Auckland*

#### S1. METHODS

##### LSD data

Full experimental details can be found in the original publication [1], but are reiterated here.

###### *Ethical statement*

This study was approved by the National Research Ethics Service (London - West London & GTAC Research Ethics Committee, REC Reference 13/LO/0229) and was conducted in accordance with the Declaration of Helsinki, the International Committee on Harmonization Good Clinical Practice guidelines, and the National Health Service Research Governance Framework. Imperial College London sponsored the research. All participants gave written consent following a comprehensive briefing and health assessment [1].

###### *Participants*

Twenty participants were recruited via word of mouth and underwent a health screening encompassing an electrocardiogram, standard blood tests, urine analysis, and a brief psychiatric interview. Exclusion encompassed age <21, a personal psychiatric diagnosis, a family history of psychotic disorders, and lack of prior exposure to classic psychedelic drugs (such as LSD, mescaline, psilocybin/magic mushrooms, or DMT/ayahuasca), any psychedelic drug intake within 6 weeks before the initial scanning session, pregnancy, excessive alcohol consumption (i.e., > 40 units per week), or any significant medical condition. Participants underwent screening at Imperial's Clinical Research Facility located on the Hammersmith Hospital campus [1].

###### *Experimental procedures*

All research activities occurred at Cardiff University Brain Research Imaging Centre. The study was a single-blind, balanced cross-over design where participants were intravenously administered a placebo on one day and 75 µg of LSD via a 10 mL solution infused over a two-minute period on another, with a minimum 14-day gap between them. Following administration, participants underwent an acclimatization phase lasting around 60 minutes. During this time, they were encouraged to relax and, at intervals, were prompted to recline with eyes shut within a simulated MRI scanner to prepare them for the potentially anxiogenic environment of the real scanning environment. MRI scanning commenced approximately 70 minutes after dosing and extended for roughly 60 minutes, encompassing structural scans, arterial spin labeling and fMRI using a 3T GE HDx system with 3D fast spoiled gradient echo scans in an

axial orientation. Post-MRI scanning, there was a break lasting approximately 35 minutes before the initiation of MEG scanning. Participants underwent four distinct conditions while under the scanner: eyes-open resting-state, eyes-closed resting-state, listening to music with eyes closed, and watching segments of a nature documentary. Each block contained seven-minutes worth of recording. Herein, we only analyse the eyes-closed resting-state data for consistency with the other studies. During MEG recordings, participants were positioned supine while pulse rate and blood oxygenation levels were consistently monitored. Participant compliance was tracked using an eye-tracking camera, and continuous monitoring of participants' head position was ensured using three fiducial coils placed at the nasion and pre-auricular points. MEG recordings were conducted using a CTF 275-channel radial gradiometer system, sampled at 1200 Hz within a band-pass range of 0–300 Hz. An additional 29 reference channels were recorded for noise cancellation, while the main sensors were analyzed as synthetic third-order gradiometers. Four out of the 275 channels were disabled due to excessive sensor noise [1].

##### *Preprocessing*

All preprocessing steps were performed using the FieldTrip toolbox. Recordings were initially high-pass filtered at 1 Hz, and segmented into 2 s epochs. Each epoch was inspected, and those with physiological artifacts (ocular, muscular, respiratory, or cardiovascular) were removed. Further, an automated rejection scheme was performed such that, if the mean spectral power in the 105 Hz to 145 Hz band of 30 predefined gradiometers at the edge of the MEG dewar exceeded 10 fT, the epoch was eliminated. ICA was then performed, and in addition to faulty components being removed manually, any component that exhibited an $r > .1$ with the EOG/EMG was removed [1]. Subsequently, a low-pass filter at 150 Hz was applied, data downsampled to 600 Hz, and line noise suppressed by subtracting a least-squares-fit sinusoidal signal at 50 Hz and harmonics. Source-localisation was done to the centroids of the AAL-90 atlas [2], which was nonlinearly inverse-warped to subject-specific grids using the subjects' structural MRI scans. Source time series were estimated with a regularized LCMV beamformer, with 5% regularisation for the whitened data covariance, aligning each source with the dominant local field orientation. Five participants were excluded from analysis due to failing to complete all scanning conditions, or because of excessive movement [3].

#### Psilocybin data

Full experimental details can be found in the original publication [4], but are reiterated here.

##### *Ethical statement*

This study was approved by the National Research Ethics Service (Bath Research Ethics Committee, REC Reference 08/H0101/144) This study was approved by the National Research Ethics Service and was conducted in accordance with the Declaration of Helsinki and the National Health Service Research Governance Framework. All participants gave written consent following a comprehensive briefing and health assessment [4].

##### *Participants*

Fifteen healthy male participants (mean age 34.5 years, SD 2.3 years) were provided informed consent and were enrolled following approval by a National Health Service research ethics committee. Participants underwent health screening encompassing an electrocardiogram, routine blood tests, urine analysis for substances of abuse, and a psychiatric interview, including disclosure of drug-taking histories. Exclusion criteria included age <21 years, pregnancy, personal or family history of psychiatric disorder (including psychotic disorders), substance dependence, cardiovascular or other significant medical conditions, claustrophobia, blood or needle phobia, and any significant adverse response to a hallucinogenic drug. All participants had prior exposure to classic psychedelic drugs but no psychedelic drug intake within 6 weeks before the study [4].

##### *Experimental procedures*

The study employed a fixed-order design where participants underwent two MEG scanning sessions on each day, one using a placebo saline infusion and one using psilocybin, with participants blinded to the infusion order, but psilocybin

(2 mg in 10 ml saline) was always administered second. Following a baseline period, the assigned solution was infused over 60 seconds, after which 5 minutes of resting MEG data were immediately recorded, with subjective ratings given of their perceived "high" on a 0–100 scale via button press. Subsequently, a 10-minute visuomotor stimulation task was performed, followed by another subjective rating. Pulse rate and blood oxygenation levels were continuously monitored throughout. Post-scanning, participants completed a 23-item rating scale addressing characteristic psychedelic effects, informed by the 5D-ASC scale but refined based on prior empirical work to 22 items plus one control item for concision. MEG recordings were conducted using a CTF 275-channel radial gradiometer system sampled at 600 Hz with a 0–150 Hz band-pass filter. An additional 29 reference channels were recorded for noise cancellation, and primary sensors were analyzed as synthetic third-order gradiometers. Three of the 275 channels were disabled due to excessive sensor noise. Participants were fitted with three electromagnetic head coils at the nasion and preauricular points, localized relative to the MEG system immediately before and after each session. Vertical and horizontal electrooculograms, an electrocardiogram, and an electromyogram from the right first dorsal interosseus were recorded alongside MEG data. Finger movements during the visuomotor task were tracked using an optical displacement system. 3D Fast Spoiled Gradient Recalled anatomical MRI scans were acquired using a 3T General Electric HDx scanner with an eight-channel receive-only head coil, with 1 mm isotropic voxel resolution in an oblique–axial orientation. [4].

##### *Preprocessing*

All preprocessing was performed using the FieldTrip toolbox. Recordings were high-pass filtered at 1 Hz to suppress slow transients and segmented into 2-second epochs. Each epoch was visually inspected, and those containing gross physiological artifacts were removed from analysis. One participant was excluded from all MEG analyses due to excessive artifacts [4]. Subsequently, a low-pass filter at 150 Hz was applied, data downsampled to 600 Hz, and line noise suppressed by subtracting a least-squares-fit sinusoidal signal at 50 Hz and harmonics. Source localization was performed to the centroids of the AAL-90 atlas [2], which was nonlinearly inverse-warped to subject-specific grids using participants' structural MRI scans. Source time series were estimated using a regularized LCMV beamformer with 5% regularization for the whitened data covariance, aligning each source with the dominant local field orientation.

#### Ketamine data

Full experimental details can be found in the original publication [5], but are reiterated here.

##### *Ethical statement*

This study was approved by the National Research Ethics Service (Wales REC 2, REC Reference 13/WA/0059) and was conducted in accordance with the Declaration of Helsinki and the National Health Service Research Governance Framework. Cardiff University sponsored the research. All participants gave written consent following a comprehensive briefing and health assessment [5].

##### *Participants*

Nineteen healthy male participants were recruited (mean age = 25.8 years, range = 19–39). Exclusion encompassed any current or previous psychiatric disorder (determined via the Mini-International Neuropsychiatric Interview), current recreational or prescription drug use, contraindications for MEG/MRI, needle phobia, age outside the 18–45 range, smoking status, American Society of Anesthesiologists Physical Status exceeding 1, and body mass index outside 18–30 kg/m$^2$ [5].

##### *Experimental procedures*

This study was a single-blind crossover study. Participants underwent two scanning sessions on separate days: one with a placebo saline infusion and one with ketamine, in a balanced design. Each session commenced with 5 minutes of resting, followed by infusion. Each participant received an initial bolus of 0.25 mg/kg delivered over approximately 1 minute, followed by maintenance infusion at a rate of 0.375 mg/kg/h. Ten minutes of resting MEG were recorded following the start of the infusion. At this point, participants were prompted to provide a rating of their subjective "high" on a scale between 0 and 40, using a two-digit button box. Measures of saccadic eye movement velocity were then taken. During MEG recordings, participants were positioned supine while pulse rate and blood oxygenation levels were consistently monitored via a probe over their left hand index finger, with the intravenous cannula inserted on the back of the left wrist. Participants wore a nasal cannula attached to medical oxygen, which was to be used in case of an emergency. MEG recordings were conducted using a CTF 275-channel radial gradiometer system, sampled at 1200 Hz within a band-pass range of 0–300 Hz. An additional 29 reference channels were recorded for noise cancellation, while the main sensors were analyzed as synthetic third-order gradiometers. Participants' ECG, horizontal and vertical electrooculograms (EOG), and electromyograms (EMG) from bilateral frontalis and temporalis muscles were recorded. All subjects had a structural MRI scan either as part of the study, or as volunteers in previous studies in Cardiff University Brain Research Imaging Centre (CUBRIC), these were 3D Fast Spoiled Gradient Recalled anatomical MRI scans using a 3T General Electric HDx scanner with an eight-channel receive-only head coil, with 1 mm isotropic voxel resolution in an oblique–axial orientation [5].

##### *Preprocessing*

All preprocessing steps were performed using the FieldTrip toolbox. Recordings were initially high-pass filtered at 1 Hz, and segmented into 2 s epochs. Each epoch was inspected, and those with physiological artifacts (ocular, muscular, respiratory, or cardiovascular) were removed. Further, an automated rejection scheme was performed such that, if the mean spectral power in the 105 Hz to 145 Hz band of 30 predefined gradiometers at the edge of the MEG dewar exceeded 10 $fT^2$, the epoch was eliminated. ICA was then performed, and in addition to faulty components being removed manually, any component that exhibited an $r > .1$ with the EOG/EMG was removed [1]. Subsequently, a low-pass filter at 150 Hz was applied, data downsampled to 600 Hz, and line noise suppressed by subtracting a least-squares-fit sinusoidal signal at 50 Hz and harmonics. Source localization was performed at the centroids of the AAL-90 atlas [2], which was nonlinearly inverse-warped to subject-specific grids using participants' structural MRI scans. Source time series were estimated with a regularized LCMV beamformer with 4 mm isotropic resolution, with 5% regularisation for the whitened data covariance, aligning each source with the dominant local field orientation [5, 6].

#### DMT data

Full experimental details can be found in the original publication [7], but are reiterated here.

##### *Ethical statement*

This study was approved by the National Research Ethics Service (London - Brent Research Ethics Committee, REC Reference 16/LO/0897), and was conducted in accordance with the revised Declaration of Helsinki, the International Committee on Harmonisation Good Clinical Practice guidelines, and the National Health Service Research Governance Framework. Imperial College London sponsored the research, which was conducted under a Home Office licence for research with Schedule 1 drugs. All participants gave written informed consent [7].

##### *Participants*

Thirteen healthy participants (6 female, 7 male, age 34.4±9.1 years) were recruited. Screening consisted of routine physical examination, electrocardiogram, blood pressure and pulse, routine blood tests, and a psychiatric interview conducted by a medic. Exclusion criteria included age <18 years, no previous experience with a psychedelic/hallucinogenic drug, personal history of diagnosed psychiatric illness, immediate family history of psychotic disorder, excessive alcohol use (>40 units per week), and blood or needle phobia. A urine test for drugs of abuse and pregnancy (where applicable), and a breathalyser test, were conducted on each study day prior to drug administration [7].

##### *Experimental procedures*

Participants attended two experimental sessions at the National Institute for Health Research (NIHR) Imperial Clinical Research Facility, resting in a semi-supine position with eyes closed throughout, aided by eye shades. EEG data were collected from one minute prior to drug or placebo administration up to twenty minutes after. Each participant received one of four doses of DMT fumarate intravenously (three received 7 mg, four received 14 mg, one received 18 mg, and five received 20 mg) in 2 mL sterile saline infused over 30 seconds, followed by a 5 mL saline flush over 15 seconds; placebo (2 mL sterile saline) followed the same procedure. A fixed-order, single-blind design was used, with placebo administered on the first visit and DMT on the second, one week later. Blood samples were collected at selected timepoints to determine plasma DMT concentration. Subjective effects were assessed via minute-to-minute intensity ratings, Visual Analogue Scales completed once acute effects had subsided, and micro-phenomenological interviews conducted the following day. EEG was recorded using a 32-channel BrainProducts system (EasycapMR 32) at a sampling rate of 1000 Hz, with a 0.1 Hz high-pass filter and 450 Hz anti-aliasing filter applied [7].

##### *Preprocessing*

EEG data were preprocessed using the FieldTrip toolbox. Data were band-pass filtered at 1 Hz to 45 Hz and visually inspected; segments containing gross artefacts were removed, as were segments during which intensity ratings were being collected. ICA was performed and components associated with eye movements and muscle activity were removed. Data were re-referenced to the average of all channels and segmented into 3 s epochs [7].

We then source localised data to the same fixed anatomical space as the other neurophysiological datasets. To do so, a mid-resolution fsaverage cortical mesh was generated by a fourth-order icosahedral subdivision (ico4), a three-layer BEM model constructed, and electrode locations were taken from the standard 10-05 template montage and registered to the fsaverage template. The forward solution leadfield matrix was then computed. To ensure consistent spatial filtering, for each subject, the data covariance matrix was estimated by pooling epochs across both the placebo and drug conditions, with rank estimated directly from the data, and an ad hoc (fixed-variance diagonal) noise covariance used for both conditions to provide a consistent neural activity index (NAI) floor. Cortical source activity was estimated via a LCMV beamformer, with 5% regularisation for the whitened data covariance, aligning each source with the dominant local field orientation. The same condition-blind spatial filter was then applied to each condition separately. Source estimates were parcellated to a surface-projected AAL-90 atlas [2], with subcortical regions excluded. Signals were extracted for each parcel by finding the dominant direction of source space normal vector orientations within each label, applying a sign-flip to time series at vertices whose orientation is more than 90°different from the dominant direction, and finally averaging across vertices at each time point within each label.

#### 5-MeO-DMT data

Full experimental details can be found in the original publication [8], but are reiterated here.

##### *Ethical statement*

This study was conducted in accordance with the Declaration of Helsinki and approved by the University College London (UCL) Research Ethics Committee (ID: 19437/004). The study was conducted in collaboration with Tandava Retreat Centre (TRC), which provided trained facilitators and facilities. All participants provided informed consent via a secure online platform after reviewing comprehensive study information. Participation was voluntary and uncompensated. Participants were informed of their right to withdraw at any stage of the study without penalty and were provided with information about sources of support in case of any distress related to the study [8].

##### *Participants*

Participants were recruited globally through advertisement on the F.I.V.E. (5-MeO-DMT Information & Vital Education) platform (www.five-meo.education), social media, news outlets, and word-of-mouth. Potential participants underwent a four-stage screening process: (1) TRC online screening, (2) TRC facilitator interview, (3) UCL online screening, and (4) UCL researcher interview (RM, GB). Exclusion criteria included: age <18 years, no prior 5-MeO-DMT experience, significant physical illness (e.g. epilepsy, heart disease), psychiatric diagnoses, use of psychiatric medications of any kind, family history of psychosis, history of adverse reactions to psychedelic drugs, and known movement under 5-MeO-DMT. Eligible participants (N = 32) were initially assigned to one of six three-day retreats, with assignments based on availability and balanced distribution. However, three participants withdrew before the retreats began. The final cohort (N = 29; 16 male, 13 female; mean age = 48.52 years, SD = 10.44, range = 34–75) was predominantly White/Caucasian (86.21%). The majority of participants held a Bachelor's degree (58.62%) and identified as non-religious (93.10%). Mean lifetime 5-MeO-DMT use was 39.03 occasions (SD = 72.91, range = 1–300) [8].

*Experimental procedures*

The study was conducted in a naturalistic setting, at a retreat centre. The study followed a three-day protocol including a preparation day consisting of orientation and participant briefing (Day 1), a dosing day for 5-MeO-DMT administration and primary data collection (Day 2), and an integration day featuring facilitator-guided group discussions (Day 3). This setup was consistent across all six retreats. Participants were instructed to refrain from consuming drugs, with the exception of nicotine containing products, for two weeks prior to dosing. On dosing day, participants fasted and abstained from caffeine. Individual dosing sessions commenced at 08:00 in 90-minute intervals. At all times during every session, two certified professional facilitators were present to ensure participant safety and provide psychological support if necessary.

Each session began with a 7-minute baseline eye-closed resting-state EEG recorded with participants seated in a self-selected comfortable position conducive to minimal movement. Participants were instructed to remain relaxed but wakeful, and to alert experimenters if they felt like they may fall asleep. Participants then moved to a centrally located padded recliner, where they assumed a semi-supine position equipped with a standardised neck pillow and opaque eye mask. Synthetic 5-MeO-DMT (12 mg) was vaporised using an argon gas piston vaporiser (203 °C to 210 °C, ∼120 s heating). Following a standardised 2–3 minute relaxation exercise, participants inhaled the vapour in a single breath and were instructed to maximise breath retention before exhalation, as per protocols established during preparation. EEG recording began at the moment of completed inhalation, and continued for 20 minutes post-inhalation. Consistent with the resting-state baseline, participants were instructed to keep eyes-closed during the entire recording. The EEG system was a saline-based 64-channel ANT Neuro Waveguard Net connected to an NA-261 EEG amplifier, recorded at 500 Hz. Ambient non-percussive music was played throughout the baseline and drug period[8].

*Preprocessing*

Preprocessing was performed using custom scripts using MNE-Python [9]. Participants were excluded from analysis if their physiological artifacts (ocular, muscular, respiratory, or cardiovascular) were substantial enough to necessitate the removal of over 50% of the data collected within the first 10 minutes post-drug administration (10 participants). Data was filtered using a finite impulse response (FIR) one-pass, zero-phase, non-causal bandpass filter at 0.1–50 Hz. Data was segmented into non-overlapping 3 s epochs. Electrolyte bridging was computed using the intrinsic Hjorth algorithm with a 16 $\mu V^2$ cut-off [10, 11]. Participants with ≥1% of potential total bridges, or significant bridging that localises such that interpolation would depend on sensors which are also bridged, were excluded from further analysis (1 participant). For those with <1%, spherical spline interpolation was performed via the generation of virtual channels between bridged electrodes (Rest: 3.684 ± 2.076, 5-MeO-DMT: 1.158 ± 0.785 bridges). Sensor time series were manually inspected to identify bad channels, participants with >20 bad channels were excluded from further analysis (0 participants), and those with <20 bad channels had these interpolated from surrounding sensors (Rest: 7.579 ± 1.049, 5-MeO-DMT: 7.368 ± 1.06 channel). Extended infomax ICA was performed with 40 components, and those manually deemed to be equipment noise or physiological artefacts were zeroed out and excluded from sensor signal reconstruction (Rest: 4.842 ± 0.766, 5-MeO-DMT: 5.053 ± 0.89 components). Epochs corrupted by noise or physiological artefacts were manually removed (Rest: 3.611 ± 2.267, 5-MeO-DMT: 38.105 ± 10.718 epochs). Data was baseline corrected and rereferenced to an average reference. This resulted in 19 subjects with epoch data [8].

We then source localised data to the same fixed anatomical space as the other neurophysiological datasets. To do so, a mid-resolution fsaverage cortical mesh was generated by a fourth-order icosahedral subdivision (ico4), a three-layer BEM model constructed, and electrode locations were taken from the standard 10-20 template montage and registered to the fsaverage template. To ensure consistent spatial filtering, for each subject, the data covariance matrix was estimated by pooling epochs across both conditions, with rank estimated directly from the data, and an ad hoc (fixed-variance diagonal) noise covariance used for both conditions. Cortical source activity was estimated via a LCMV beamformer, with 5% regularisation for the whitened data covariance, aligning each source with the dominant local field orientation. The same condition-blind spatial filter was then applied to each condition separately. Source estimates were parcellated to a surface-projected AAL-90 atlas [2], with subcortical regions excluded. Signals were extracted for each parcel by finding the dominant direction of source space normal vector orientations within each label, applying a sign-flip to time series at vertices whose orientation is more than 90°different from the dominant direction, and finally averaging across vertices at each time point within each label.

#### S2. RESULTS

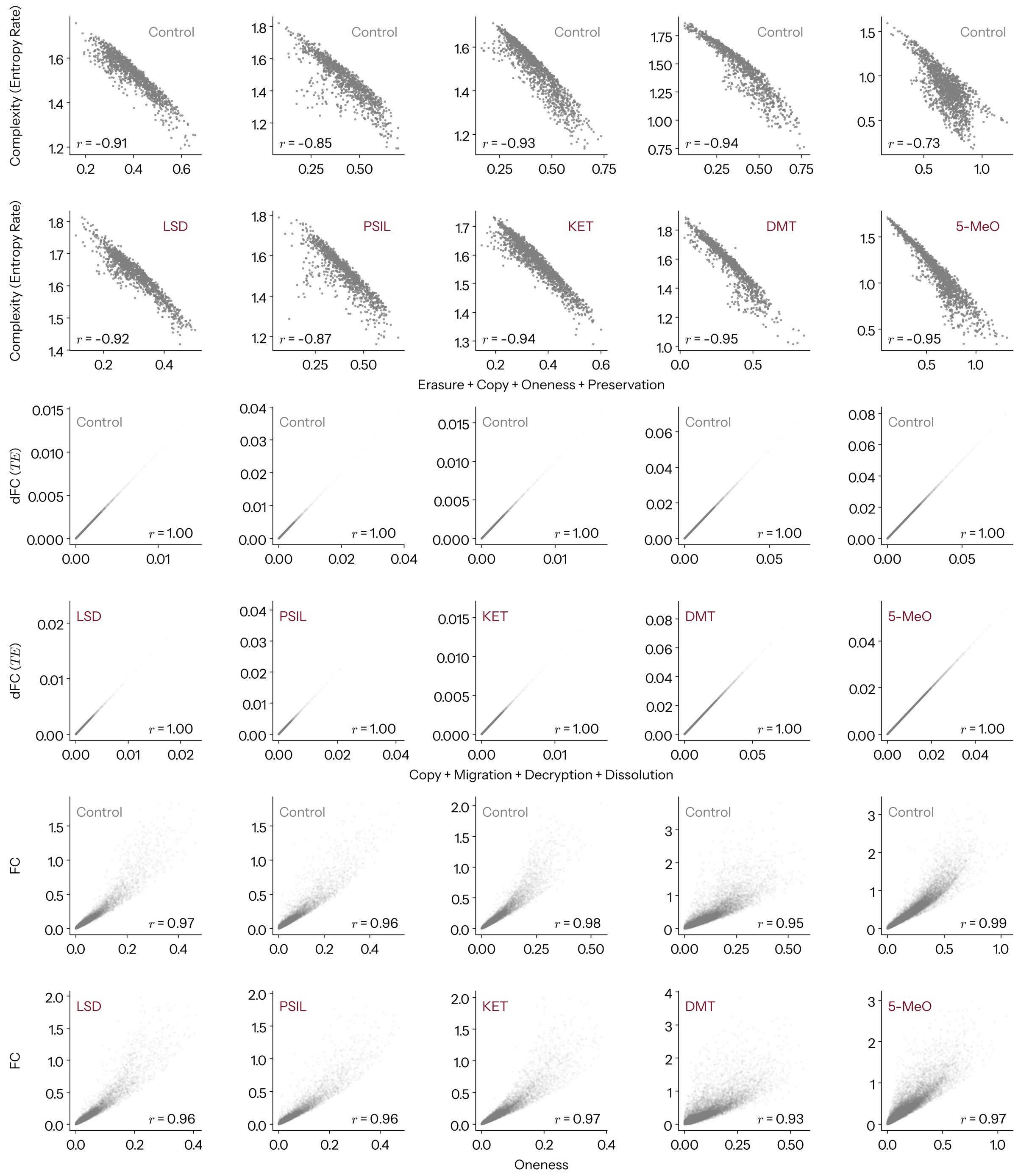


FIG. S1. **Accounting for existing measures with information dynamics.** Complement to main Fig. 2 showing empirical relationships between measures and their (aggregate) information dynamical constituents, independently for each condition. Upper block Complexity via State-Space Entropy rate versus Active Information Storage (AIS), which is the sum of Erasure, Copy, Oneness, and Preservation. Middle block Directed Functional Connectivity via Transfer entropy, versus Copy, Migration, Decryption and Dissolution. Lower block Functional Connectivity versus Oneness. Top row plots are for respective control conditions, and the bottom rows for each respective drug.

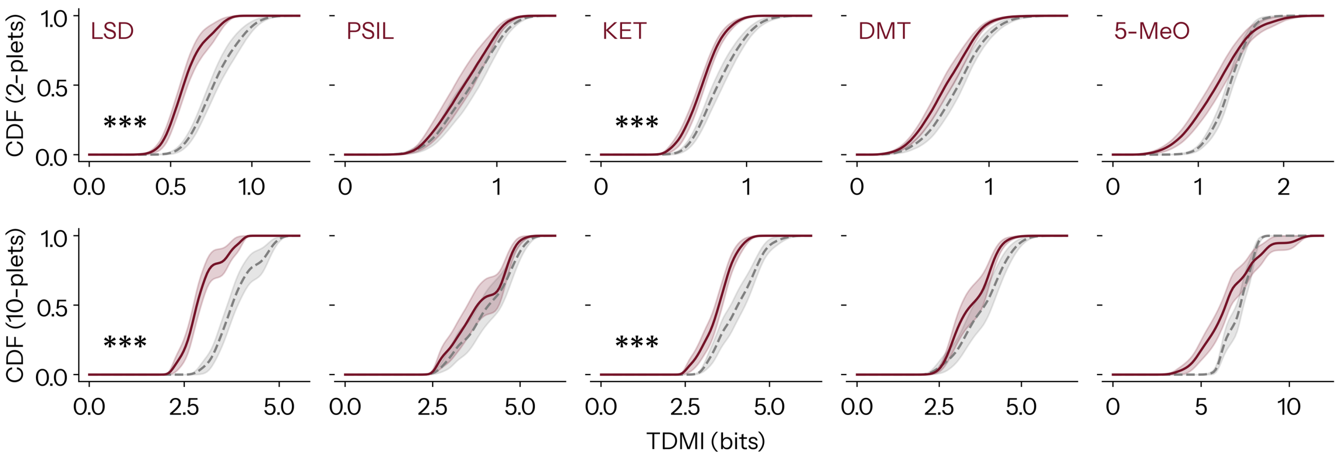


FIG. S2. **Psychedelics alter the TDMI.** Psychedelics reduce the total amount of information flowing from past to future (TDMI, $I(\mathbf{X}_t; \mathbf{X}_{t'})$), suggesting that alterations in absolute unnormalised modes of information dynamics may trivially reflect alterations in this quantity. Effect is consistent across 2-plets (top) and 10-plets (bottom). (Grey dashed line for control condition, purple for drug - Mean ± SEM.)

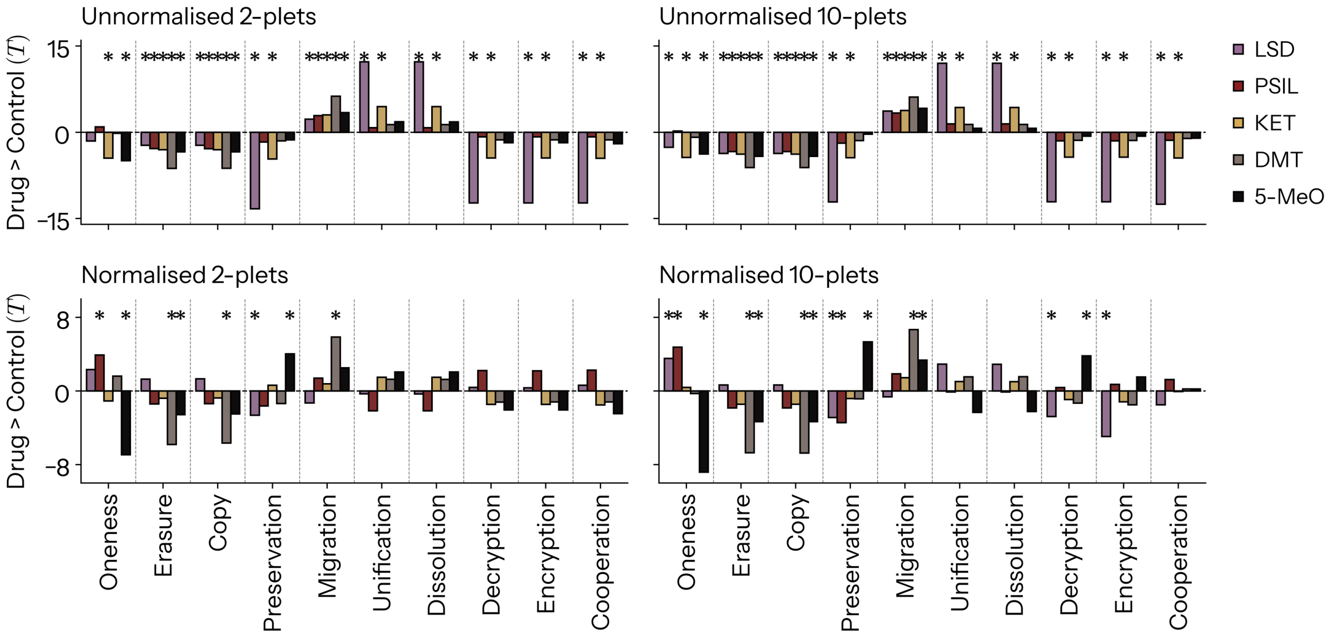


FIG. S3. **Effect of psychedelics on global information dynamics** $T$-statistic plots for global (mean) alterations in modes of information dynamics for each drug, for both 2-plets and 10-plets, as well as unnormalised (absolute) versus normalised (relative). Results are consistent across orders of interaction, with 10-plets possessing stronger statistical power. Results markedly differ when normalised.

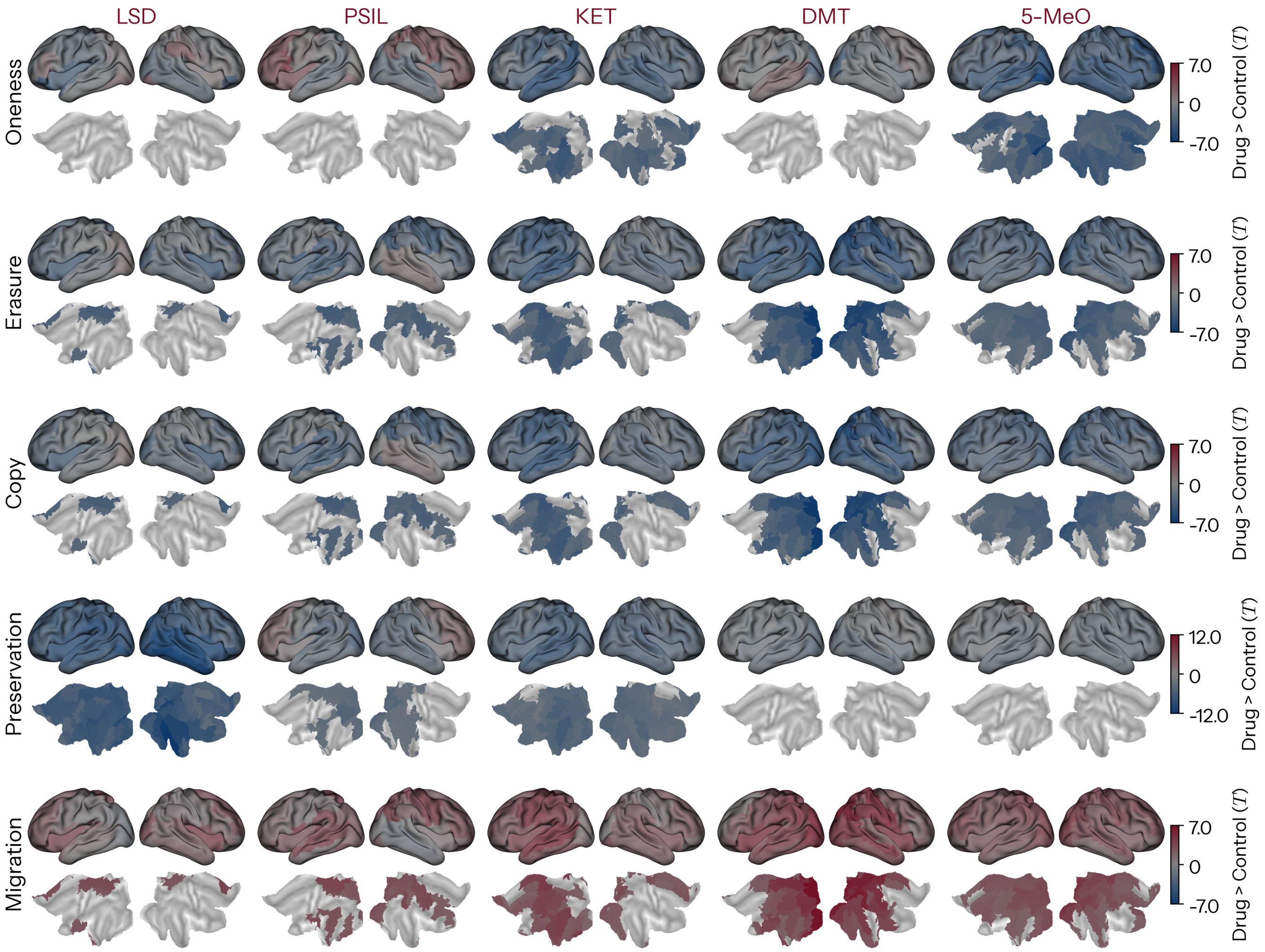


FIG. S4. **Effect of psychedelics on absolute regional lower-order information dynamics** Topographic statistical contrast maps for unnormalised Oneness, Erasure, Copy, Preservation, and Migration for each drug. Inflated surface for $T$-statistic (upper), flat map thresholded at $p_{\mathrm{TFCE}} < .05$.

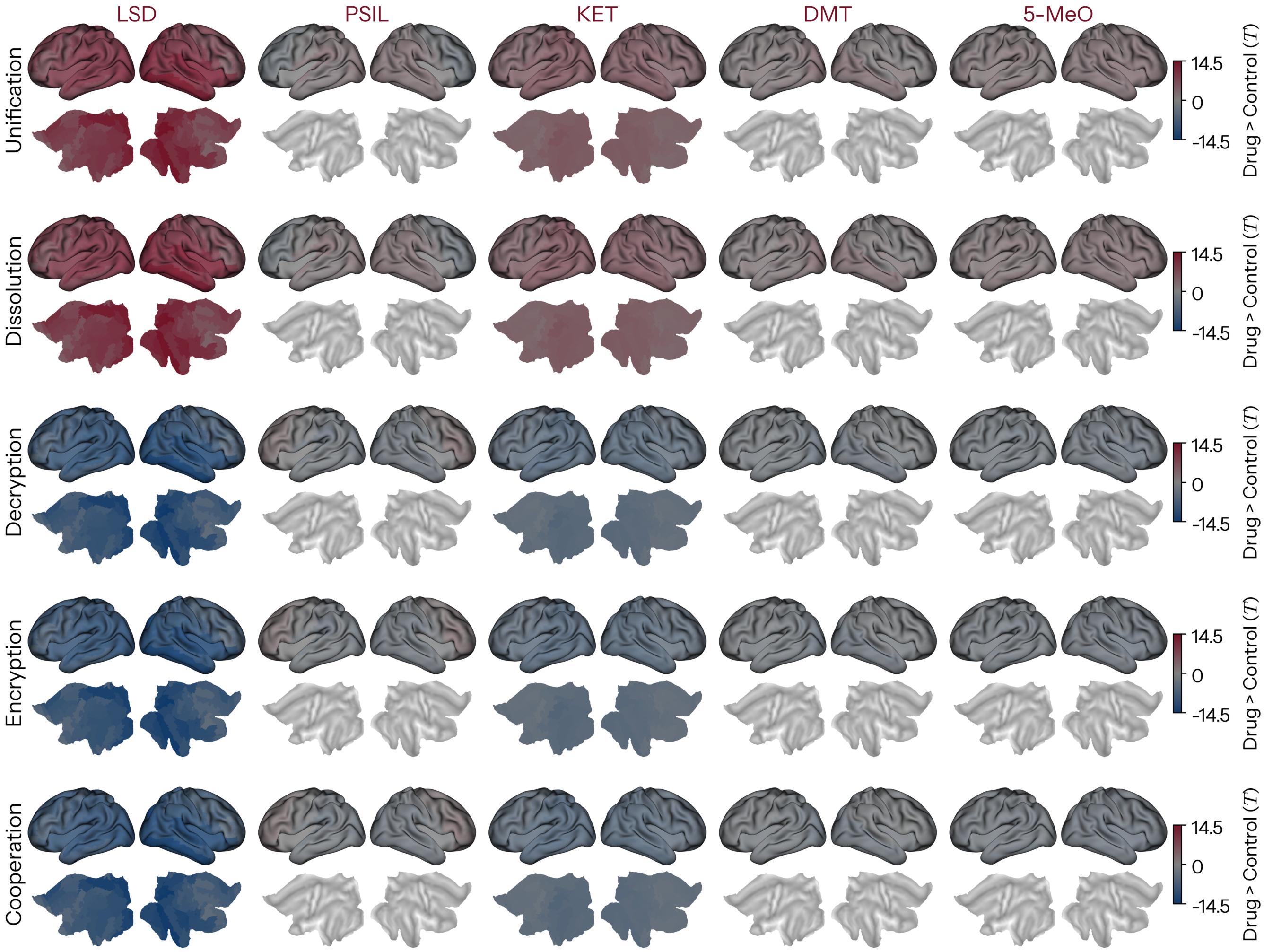


FIG. S5. **Effect of psychedelics on absolute regional higher-order information dynamics** Topographic statistical contrast maps for unnormalised Unification, Dissolution, Decryption, Encryption, and Cooperation for each drug. Inflated surface for raw $T$-statistic (upper), flat map thresholded at $p_{\mathrm{TFCE}} < .05$.

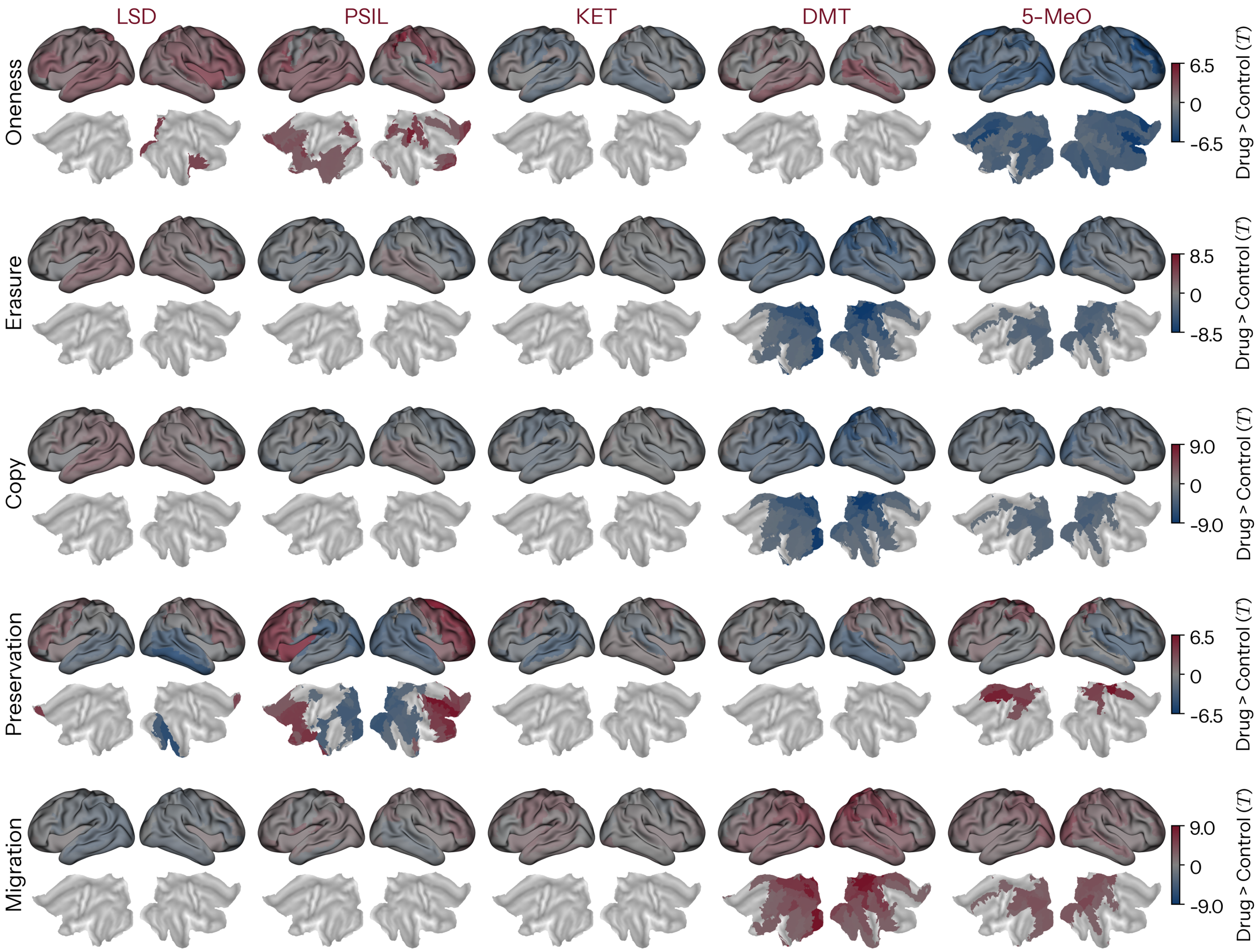


FIG. S6. **Effect of psychedelics on relative regional lower-order information dynamics** Topographic statistical contrast maps for normalised Oneness, Erasure, Copy, Preservation, and Migration for each drug. Inflated surface for $T$-statistic (upper), flat map thresholded at $p_{\text{TFCE}} < .05$.

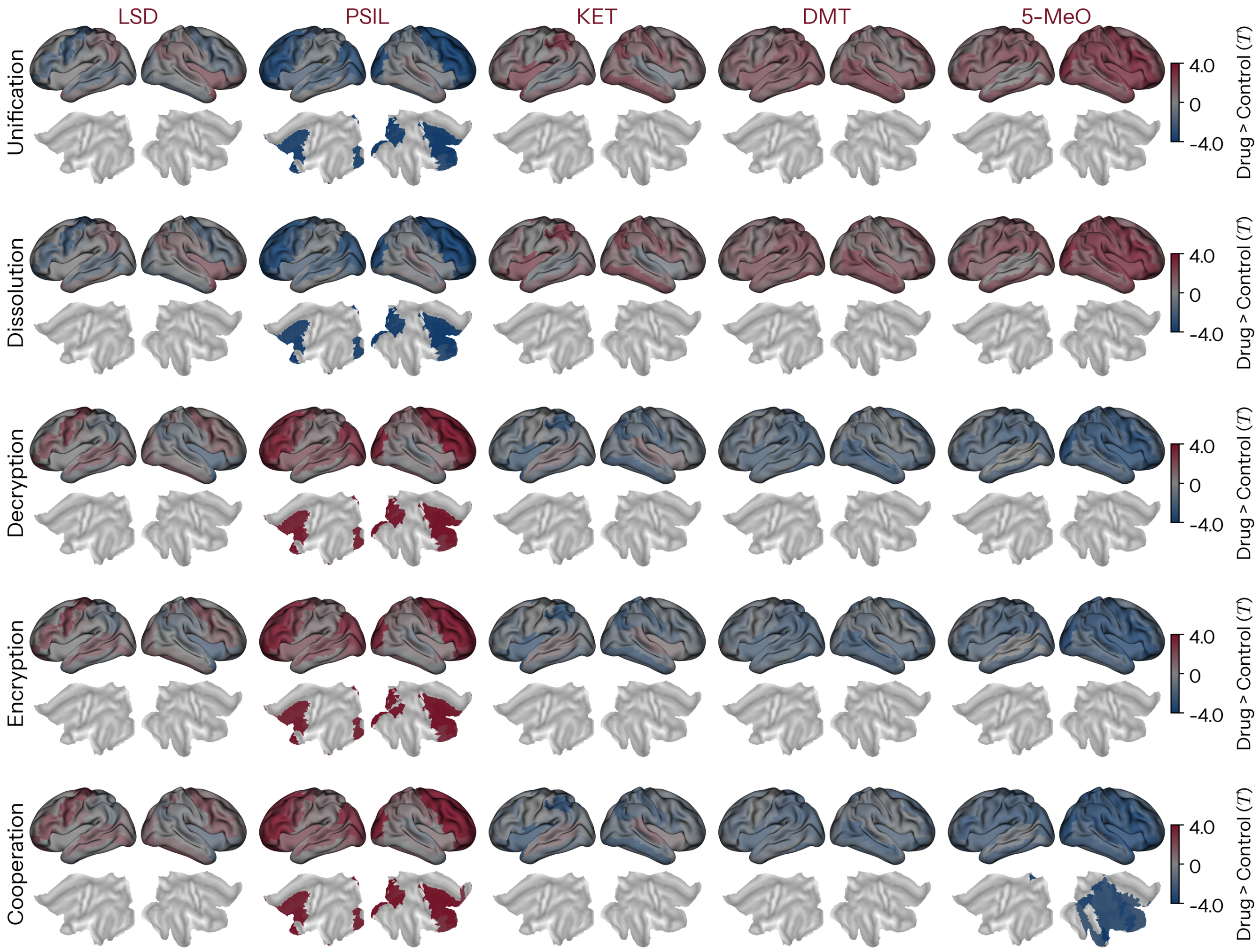


FIG. S7. **Effect of psychedelics on relative regional higher-order information dynamics** Topographic statistical contrast maps for normalised Unification, Dissolution, Decryption, Encryption, and Cooperation for each drug. Inflated surface for $T$-statistic (upper), flat map thresholded at $p_{\mathrm{TFCE}} < .05$.

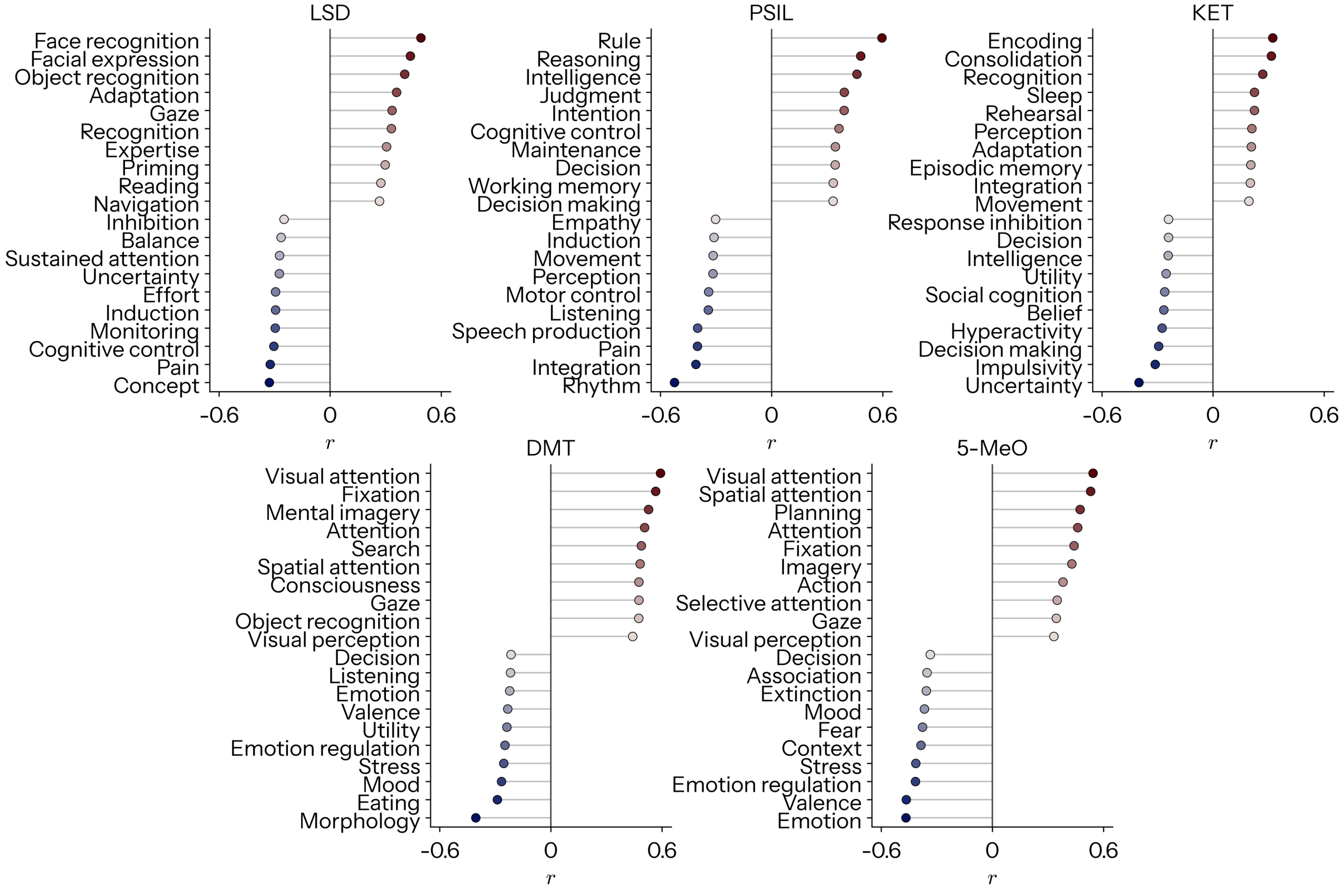


FIG. S8. **Neurosynth decoding of meta-analytic information dynamics topographies for each drug** Correlation of fully pooled meta-analytic maps for each drug with meta-analytic cognitive maps from Neurosynth. The top ten most correlated and anti-correlated terms are displayed.

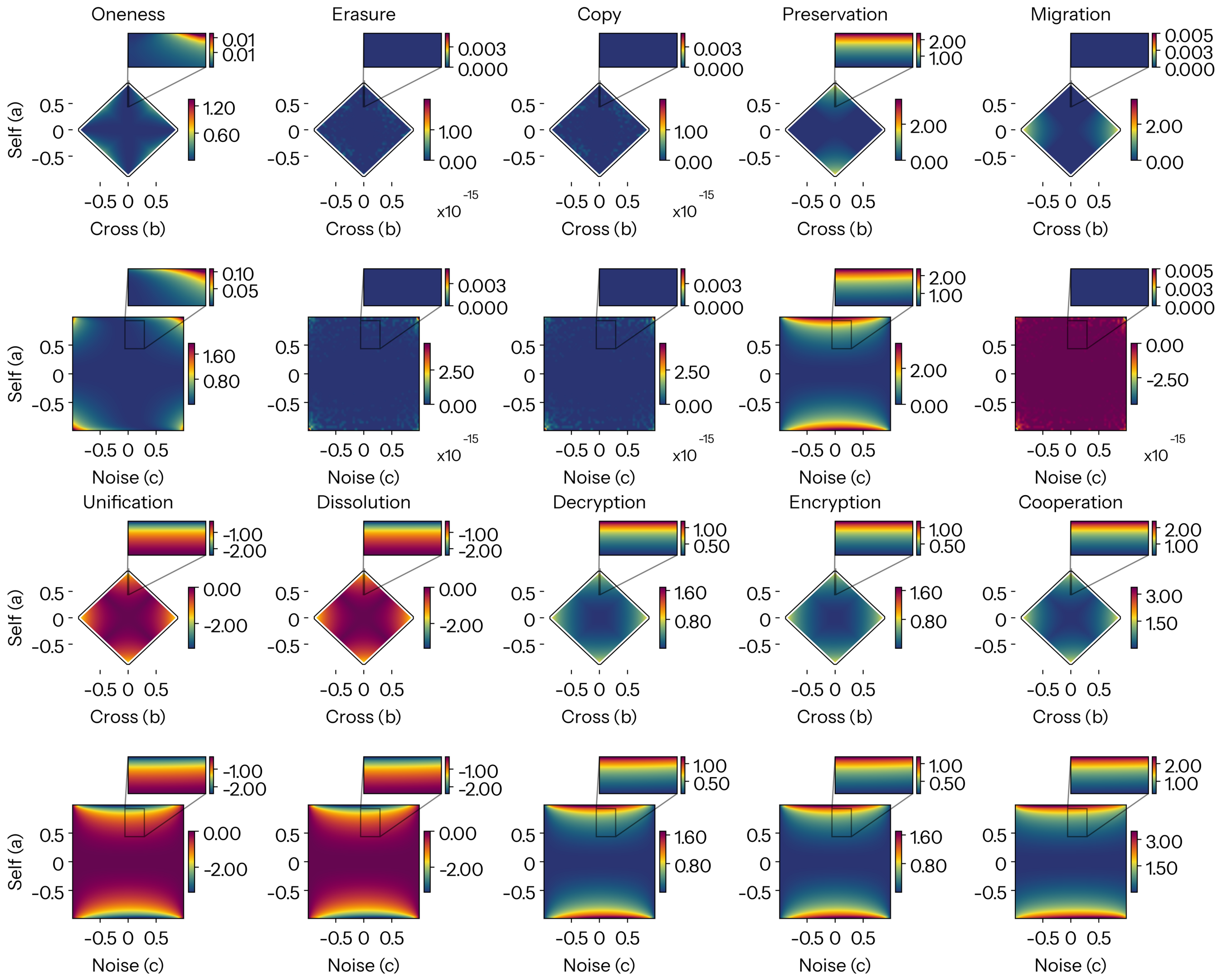


FIG. S9. **Reconfigurations in absolute information dynamics in the sweep of model parameters.** First row: $a$ versus $c$ with fixed $b = 0.02$. Second row: $a$ versus $b$ with fixed $c = 0.06$. $a$ vs $b$ plot has black borders denoting the boundary of model parameters that yield valid covariance matrices. Zoomed inlets display the space occupied by empirical coefficients.

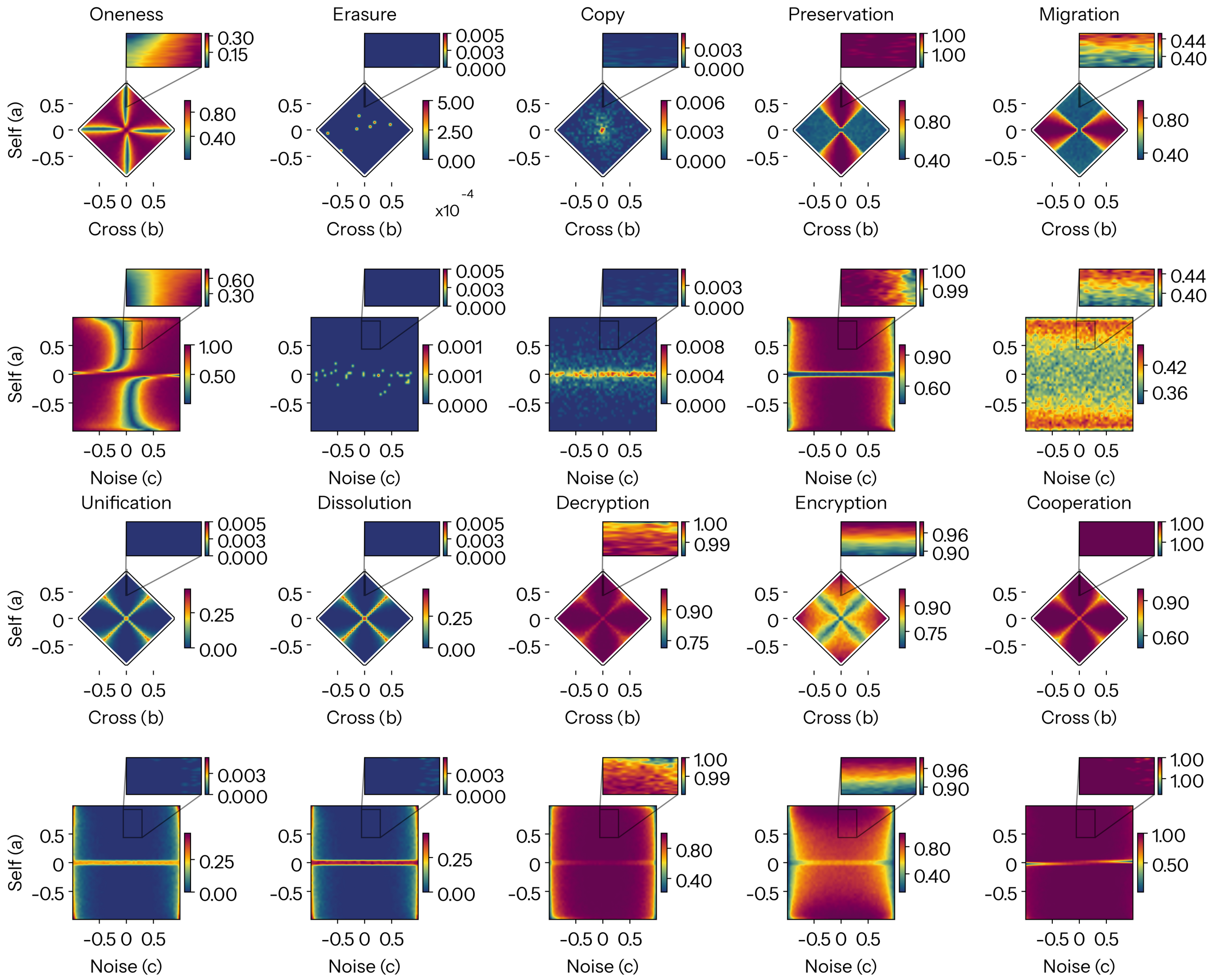


FIG. S10. **Reconfigurations in relative information dynamics in the sweep of model parameters.** First row: $a$ versus $c$ with fixed $b = 0.02$. Second row: $a$ versus $b$ with fixed $c = 0.06$. $a$ vs $b$ plot has black borders denoting the boundary of model parameters that yield valid covariance matrices. Zoomed inlets display the space occupied by empirical coefficients.

TABLE S1. **NeuroSynth terms.** 123 terms that overlap between the NeuroSynth [12] and the Cognitive Atlas [13]

| | | | |
|---|---|---|---|
| action | adaptation | addiction | anticipation |
| anxiety | arousal | association | attention |
| autobiographical memory | balance | belief | categorization |
| cognitive control | communication | competition | concept |
| consciousness | consolidation | context | coordination |
| decision | decision making | detection | discrimination |
| distraction | eating | efficiency | effort |
| emotion | emotion regulation | empathy | encoding |
| episodic memory | expectancy | expertise | extinction |
| face recognition | facial expression | familiarity | fear |
| fixation | focus | gaze | goal |
| hyperactivity | imagery | impulsivity | induction |
| inference | inhibition | insight | integration |
| intelligence | intention | interference | judgment |
| knowledge | language | language comprehension | learning |
| listening | localization | loss | maintenance |
| manipulation | meaning | memory | memory retrieval |
| mental imagery | monitoring | mood | morphology |
| motor control | movement | multisensory | naming |
| navigation | object recognition | pain | perception |
| planning | priming | psychosis | reading |
| reasoning | recall | recognition | rehearsal |
| reinforcement learning | response inhibition | response selection | retention |
| retrieval | reward anticipation | rhythm | risk |
| rule | salience | search | selective attention |
| semantic memory | sentence comprehension | skill | sleep |
| social cognition | spatial attention | speech perception | speech production |
| strategy | strength | stress | sustained attention |
| task difficulty | thought | uncertainty | updating |
| utility | valence | verbal fluency | visual attention |
| visual perception | word recognition | working memory | |

TABLE S2: **Statistics for average fitted model coefficients.**

| Drug | Parameter | $T$ | $d$ | $BF_{10}$ | $p$ | $p_{FDR}$ | DoF |
|---|---|---|---|---|---|---|---|
| LSD | $a$ | −12.944 | 1.619 | $3.01 \times 10^6$ | $3.52 \times 10^{-9}$ | $1.76 \times 10^{-8}$ | 14 |
| LSD | $b$ | 0.931 | 0.330 | 0.381 | 0.367 | 0.757 | 14 |
| LSD | $c$ | 1.410 | 0.407 | 0.599 | 0.180 | 0.301 | 14 |
| LSD | $\rho$ | −13.367 | 1.801 | $4.42 \times 10^6$ | $2.31 \times 10^{-9}$ | $1.16 \times 10^{-8}$ | 14 |
| PSIL | $a$ | −1.209 | 0.213 | 0.498 | 0.248 | 0.248 | 13 |
| PSIL | $b$ | −0.476 | 0.100 | 0.298 | 0.642 | 0.802 | 13 |
| PSIL | $c$ | 0.522 | 0.076 | 0.304 | 0.611 | 0.763 | 13 |
| PSIL | $\rho$ | −1.888 | 0.332 | 1.087 | 0.082 | 0.136 | 13 |
| KET | $a$ | −4.485 | 1.031 | $1.10 \times 10^2$ | $2.86 \times 10^{-4}$ | $7.16 \times 10^{-4}$ | 18 |
| KET | $b$ | −0.764 | 0.280 | 0.308 | 0.454 | 0.757 | 18 |
| KET | $c$ | −0.167 | 0.049 | 0.241 | 0.869 | 0.869 | 18 |
| KET | $\rho$ | −4.515 | 0.974 | $1.17 \times 10^2$ | $2.68 \times 10^{-4}$ | $6.70 \times 10^{-4}$ | 18 |
| DMT | $a$ | −1.435 | 0.602 | 0.641 | 0.177 | 0.221 | 12 |
| DMT | $b$ | −1.028 | 0.285 | 0.434 | 0.324 | 0.757 | 12 |
| DMT | $c$ | 2.041 | 0.335 | 1.339 | 0.064 | 0.160 | 12 |
| DMT | $\rho$ | −1.227 | 0.471 | 0.519 | 0.243 | 0.243 | 12 |
| 5-MeO | $a$ | −2.318 | 0.756 | 1.993 | 0.032 | 0.054 | 18 |
| 5-MeO | $b$ | −0.055 | 0.016 | 0.238 | 0.957 | 0.957 | 18 |
| 5-MeO | $c$ | −2.240 | 0.212 | 1.758 | 0.038 | 0.160 | 18 |
| 5-MeO | $\rho$ | −1.518 | 0.510 | 0.632 | 0.146 | 0.183 | 18 |
